# Neural fiber orientation across spinal cord gray matter

**DOI:** 10.64898/2026.08.06.743226

**Authors:** Sigurd Fyhn Sørensen, Jaspreet Kaur, Rune W. Berg

## Abstract

Understanding connectivity within the nervous system is central to understanding its function, and the network architecture of many brain regions has been mapped in fine detail. The sub-tract organization of neuronal projections within the human cord, particularly in gray matter is essentially unknown. Since full connectomic reconstruction is currently intractable at this scale, a tractable first step is to statistically characterize neurite orientations across spinal regions. Here we combine a ∼400 hour 9.4T ex vivo high-angular-resolution diffusion-weighted MRI spanning all segments (C1–S5) of a post-mortem human spinal cord with structure-tensor analysis of SMI-32 neurofilament fluorescence microscopy to map neurite orientation across the cord. We find that projection geometry varies systematically along the rostro-caudal axis. Cervical, lumbar, and sacral gray matter exhibit low anisotropy, high orientation dispersion, and predominantly transverse fiber populations (>91% of orientations), consistent with dense segment-local circuitry. Thoracic gray matter is categorically distinct from the rest of the cord with elevated axial diffusivity, white-matter-like anisotropy, and a fiber orientation distribution in which 60% of orientations run within a polar angle of 85–95° i.e., along the the cord’s rostro-caudal axis. Micron-resolution microscopy independently supports this longitudinal architecture. Together, these findings reveal a regional dissociation in spinal gray-matter wiring and suggest a coupling between projection geometry and the distinct processing demands of each segment.

## 1 Introduction

Neuronal projections have long been studied as a principled window into the information-passing pathways that organize nervous systems, an interest that spans a wide range of scales. At one end, it motivates understanding how pathological alterations to microstructure such as injury (1), demyelination (2, 3), beading (4), and changes in axon caliber and packing (5) can accompany, and may drive, neurological disease, a concern that has long motivated quantitative mapping of axonal microstructure. A complementary motivation treats projection anatomy as the wiring diagram, the signal-propagation architecture, underlying the computations that nervous systems require to produce behavior. Recent years have also mapped neuronal signaling to propagate through gaseous messengers, acting irrespective of synaptic apposition or axonal direction. Thus, violating a point to point connectionist picture, as anatomical connectivity does not exhaust the information passing pathways (6–8). Nevertheless, axons and neurites remain the primary anatomical substrate in our account of neuronal information transfer, and mapping their organization remains a central aim of systems neuroscience (9–11). This has been pursued, e.g., in the Human Connectome Project and comparable efforts have focused on cerebral white-matter architecture (12). Although much attention has been paid to mapping brain networks, internal connectivity in the spinal cord, besides reflex archs and the sensory motor communication projections, has received less attention. It is well known that the spinal cord has the capacity to independently generate a wide array of complicated motor functions. Graham Brown and Sherrington showed that coordinated, rhythmic motor output can be produced by the isolated cord, laying the foundation for the central pattern generator (CPG) concept, in which spinal interneuronal networks autonomously generate the timing and coordination of locomotion, scratching, and related behaviors even in the absence of descending or peripheral input (13, 14). The evidence for autonomous rhythmogenesis extends to the human spinal cord after complete injury (15) and contemporary work has addressed this capacity quantitatively and mechanistically through both *in vivo* preparations and *in silico*. Population recordings from the lumbar cord revealing that motor rhythms arise as low-dimensional rotational dynamics (16,17), which can be reproduced in recurrently connected network models that requires neither a dedicated pacemaker nor modular CPG layers (18). Such a network can be realized through a rosto-caudal spatial organization of neurite projection patterns, (19–21) highlighting why the spatial organization of orientation profiles across the spinal cord is of particular interest. With the spinal cord being computational structure in its own right rather than a passageway, and its internal wiring being crucial in facilitating its function, it then warrants the same fine-grained anatomical scrutiny that has been lavished on the brain. The human spinal cord remains comparatively uncharted, with post-mortem human microstructure and its function mapped in only a handful of efforts (5, 22–25), and a systematic account of how gray matter neuronal projections are organized beyond the the demarcation of the major rostral-caudal white-matter fiber tracts is still lacking. It is this gap, the sub-tract organization of projections in the *postmortem human* spinal cord that our work aims to address, combining ex vivo high angular resolution diffusion-weighted MRI with structure-tensor analysis of fluorescence-microscopy to map human microstructural orientation across the cord’s regions.

## 2 Results

The full experimental setup from data collection, pre-processing analysis methods and parameter testing plus selection can be found in the supplmentary methods. In this section we introduce the analysis from a macroscopic viewpoint with a focus on the scientific results. MRI data were initilialy collected and analyzed first before subsequently slicing the tissue and performing fluorescence microscopy analysis to strengthen the neurite microstructure organization discovered during our MRI analysis.

### 2.1 dwMRI: Fiber orientation in Gray and White Matter

This study was performed on a full-length skull stripped spinal cord from a deceased individual who had bequeathed her body to science and education at the Department of Cellular and Molecular Medicine (ICMM) of Copenhagen University according to Danish legislation (Health Law No. 546, Section 188). The study was approved by the head of the Body Donation Program at ICMM, ensuring ethical approval and the spinal cord originated from a 91-year-old Caucasian female without any known neurological or psychiatric diseases. It was obtained, dissected, and fixed within 24 hours post-mortem, in order to preserve tissue morphology, an immersion fixation protocol was performed using a paraformaldehyde (4%) buffer with a pH of 7.4 (26, 27). The tissue was fixed in the buffer for 2 weeks at low temperatures. For long-term storage, it was transferred to a phosphate-buffered saline solution with a pH value of 7.4. Images were acquired on a 9.4 T preclinical MRI system (BioSpec 94/30; Bruker Biospin, Ettlingen, Germany) equipped with a 1.5 T/m gradient coil. Prior to imaging, the spinal cord was placed in a plexiglas tube and immersed in fluorinert (FC-40, Sigma-Aldrich), reducing background signal Our protocol involved immersing the ex vivo tissue in perfluorocarbon prior to scanning. As perfluorocarbons contain no hydrogen protons they produce negligible background signal for conventional hydrogen-1 MRI. Producing enhanced contrast between tissue and background, increasing SNR and CNR (28). Perfluorocarbons furthermore have magnetic susceptibility close to that of tissue, reducing susceptibility artifacts at tissue boundaries, in adition to being a dense and inert fluid therby providing physical stabilization and mitigation of tissue heating (29, 30). A limited field of view of 1.6 cm relative to the length of the spinal cord 40 cm, necessitated imaging being done in 29 sections, known as Multiple Overlapping Thin Slab Acquisition (MOTSA) MRI (31, 32). Between each section-scan the spinal cord was advanced 1.4 cm by a custom-built mechanical stepper, resulting in a 0.2-cm overlap between neighboring sections. Structural MRI were acquired for each section with a T2-weighted 2D RARE sequence with associated scanning parameters, echo time (TE) = 30ms, repetition time (TR) = 7000ms, a field of view of 1.92 × 1.92 × 1.6 cm^3^, and a matrix size of 384 × 384 × 80, resulting in 50 × 50 µm^2^ in-plane resolution and a slice thickness of 200 µm, producing an anisotropic voxel of size 5e5 µm^3^. The anatomical T2 images were generated using 20 excitations, and the dwMRI was acquired for each of the 29 sections with a spin-echo high angular resolution diffusion imaging (HARDI) protocol with a single shell, b-value = 4000s/mm^2^, along with eighty motion-probing gradients encoding the 80 different diffusion directions. The total scan time totaled 400 hours of scanning time. The limited scanner field of view relative to the full cord length necessitated imaging in 29 overlapping slabs, which were subsequently stitched into a single volume spanning segments C1–S5. Preprocessing included MP-PCA denoising, eddy-current correction, Gibbs ringing suppression, and bias field correction, followed by gray and white matter segmentation using a fine-tuned convolutional neural network (Dice score_GM_ = 0.975, Dice score_WM_ = 0.987; see Supplementary S. 8) for training procedure, data augmentation pipeline and model scores. Providing a reliable anatomical tissue segmentation for all downstream analyses. The dwMRI analysis are presented in the same order as analysis took place, which also coincides with increasing model complexity, starting with the classical diffusion tensor imaging (DTI) analysis to the multi-compartment NODDI model and finishing off with high angular resolution constrained spherical deconvolution (CSD).

#### DTI reveals rostro-caudal gradients in diffusivity

To characterize the macrostructural diffusion landscape of the full spinal cord, we first applied DTI (33–35) to the high-resolution ex vivo dwMRI data spanning segments C1–S5 (Fig. 1). For analysis, the spinal cord was divided into four regions, cervical (C1–C8), thoracic (T1–T12), lumbar (L1–L5), and sacral (S1–S5), constituting 24.1%, 56.5%, 12.7% and 6.7% of the spinal cords length each, a full table of segmental length percentages can be found in (36). The sagittal slice (Fig. 1B) illustrates the full length of the cord with spinal segment boundaries demarcated, allowing metrics to be interpreted in their segmental context. Rostro-caudal profiles of mean diffusivity (MD), axial diffusivity (AD), and radial diffusivity (RD) are shown separately for gray matter (GM) and white matter (WM) in Fig. 1C. In WM, AD was consistently the largest diffusion metric and remained relatively stable across cervical, thoracic, and lumbar segments before dropping sharply in the sacral region. RD in WM was substantially lower than AD throughout, consistent with the well-established anisotropic restriction imposed by myelinated axon bundles running in the rostro-caudal direction. GM diffusivities were uniformly lower than their WM counterparts and showed a more compressed dynamic range, reflecting the greater structural complexity and multi-directional fiber architecture of gray matter. All three DTI metrics exhibited a pronounced reduction in the sacral region for both tissue types. Notably, did AD thoracic gray matter exhibit drastic increases in AD when contrasted to the remaining gray matter segments, following a quadratic curvature that peaks around segment T6. At which point the gray matter AD levels are on par with the highly parallel white matter fiber tracts. Suggesting structural reorganization across the rostral-caudal length of the cord that is detectable even at the coarse level of DTI. These segment-level differences, particularly the divergence of thoracic and sacral diffusion profiles from cervical and lumbar values, motivated the application of more sensitive, bio-physically specific models capable of resolving finer micro-structural features.

**Figure 1:**
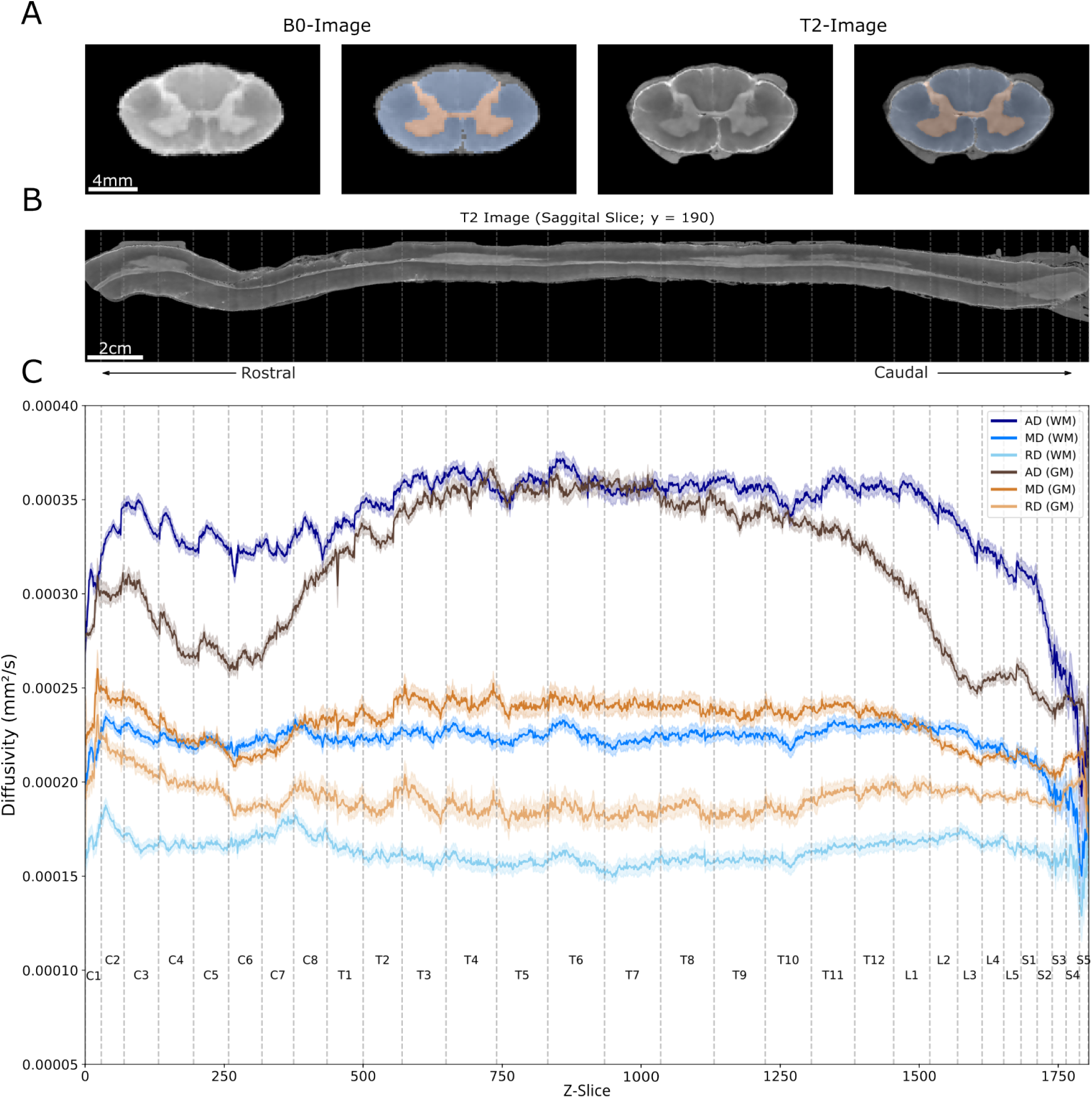
**Fig A**. Axial representation of a T2-image and B0-image overlaid with generated tissue mask. **Fig B.** T2 sagittal slice, stippled lines demarcates spinal segment boundaries. **Fig C.** Rostro-caudal gradients of DTI metrics, mean diffusivity (MD), axial diffusivity (AD), and radial diffusivity (RD) all on the scale of (mm^2^/s) and stratified by tissue type.

#### FA and ODI capture complementary micro-structural variation across segments and tissue types

To disentangle the contributions towards the observed diffusion gradients in (Fig. 1), we computed fractional anisotropy (FA) from the DTI model and orientation dispersion index (ODI) from the Neurite Orientation Disribution Density Imaging (NODDI) (37) model across all spinal segments and tissue compartments (Fig. 2). The sagittal T2 image (Fig. 2A) provides the anatomical reference for the rostro-caudal FA & ODI profiles in panel B. FA profiles (Fig. 2B, upper trace) follows similair patterns to that of AD. WM FA was consistently higher than GM FA throughout the cord, with WM & GM values peaking in the thoracic region before declining sharply through the lumbar and sacral segments. The explanation behind the similarities between AD and FA can be found in its parameterization, with FA being a metric combining all of the diffusion metrics from (Fig. 1). Minimal changes in MD and RD consequently results in FA changes being largely driven by alterations in AD values, see supplementary methods for detailed walk through of the different models and associated variables. Notably this means that the gray matter thoracic segments have significantly higher diffusion along its principal direction, with minimal changes to non-dominant orientations, thus leading to anisotropic diffusion, indicative of parallel fibers bundles. With the highest degree of gray matter fractional anisotropy being observed from T5-T8. DTI is a relatively simplistic model, being a single compartment model reducing our measured 80 diffusion direction into 3 three orthogonal eigenvectors. We therefore utilized a multi compartment model, namely NODDI from which we can derive more accurate measures of neurite orientations profiles. The ODI profiles (Fig. 2B, lower trace) showed an inverse pattern to that of FA, with lower ODI values are indicative of coherent fiber orientations. GM ODI was markedly higher than WM ODI at cervical, lumbar and sacral segments, with the thoracic GM showed a pronounced local minimum in ODI relative to adjacent segments, tangent to WM ODI levels for T4-T10. Supporting the notion that thoracic GM neurites are more coherently aligned than those of any other gray matter region. To a degree that makes GM assimilate coherent longitudinal WM fiber bundles. Probability density distributions stratified by segment and tissue type (Fig. 2C–D) confirmed that thoracic GM occupies a distinct position in both FA and ODI space. Its FA distribution (Fig. 2C) is shifted to higher values relative to cervical and lumbar GM, with a thoracic x̃ = 0.35, µ = 0.36 and σ = 0.08, compared with cervical GM x̃ = 0.20, µ = 0.22 and σ = 0.09. Correspondingly, the thoracic GM ODI distribution (Fig. 2D) is markedly narrower and shifted toward lower values, with a x̃ = 0.39, µ = 0.40 and σ = 0.06, substantially below the µ ∈ [0.6, 0.75] observed in cervical, lumbar and sacral GM. The combination of higher anisotropy paired with a lower orientation dispersion from the multi-compartment model, is a clear signature of a more coherent, directionally organized neurite population in thoracic gray matter. Pairwise effect size heatmaps (Cohen’s d; Fig. 2E–F) quantified the magnitude of these differences. For FA (Fig. 2E), thoracic GM showed a large positive effect size relative to cervical GM (d = 1.62), lumbar GM (d = 1.95), and sacral GM (d = 2.45), indicating substantially higher FA in thoracic gray matter. For ODI (Fig. 2F), the corresponding comparisons revealed large negative effect sizes: thoracic GM versus cervical GM (d = −1.93), thoracic GM versus lumbar GM (d = −2.38), and thoracic GM versus sacral GM (d = −3.30), the largest pairwise difference observed across all ODI comparisons in the dataset. See (S. 9) for a comparison of all segment x tissue combinatorics. These large effect sizes were confirmed as statistically significant under a linear mixed model accounting for spatial autocorrelation, interaction effects and corrected for multiple comparisons using Tukey’s HSD (S. 10). Together, the FA and ODI results converge on a consistent picture, thoracic gray matter harbors a markedly more coherent and directionally organized neurite population than any other gray matter region of the spinal cord.

**Figure 2:**
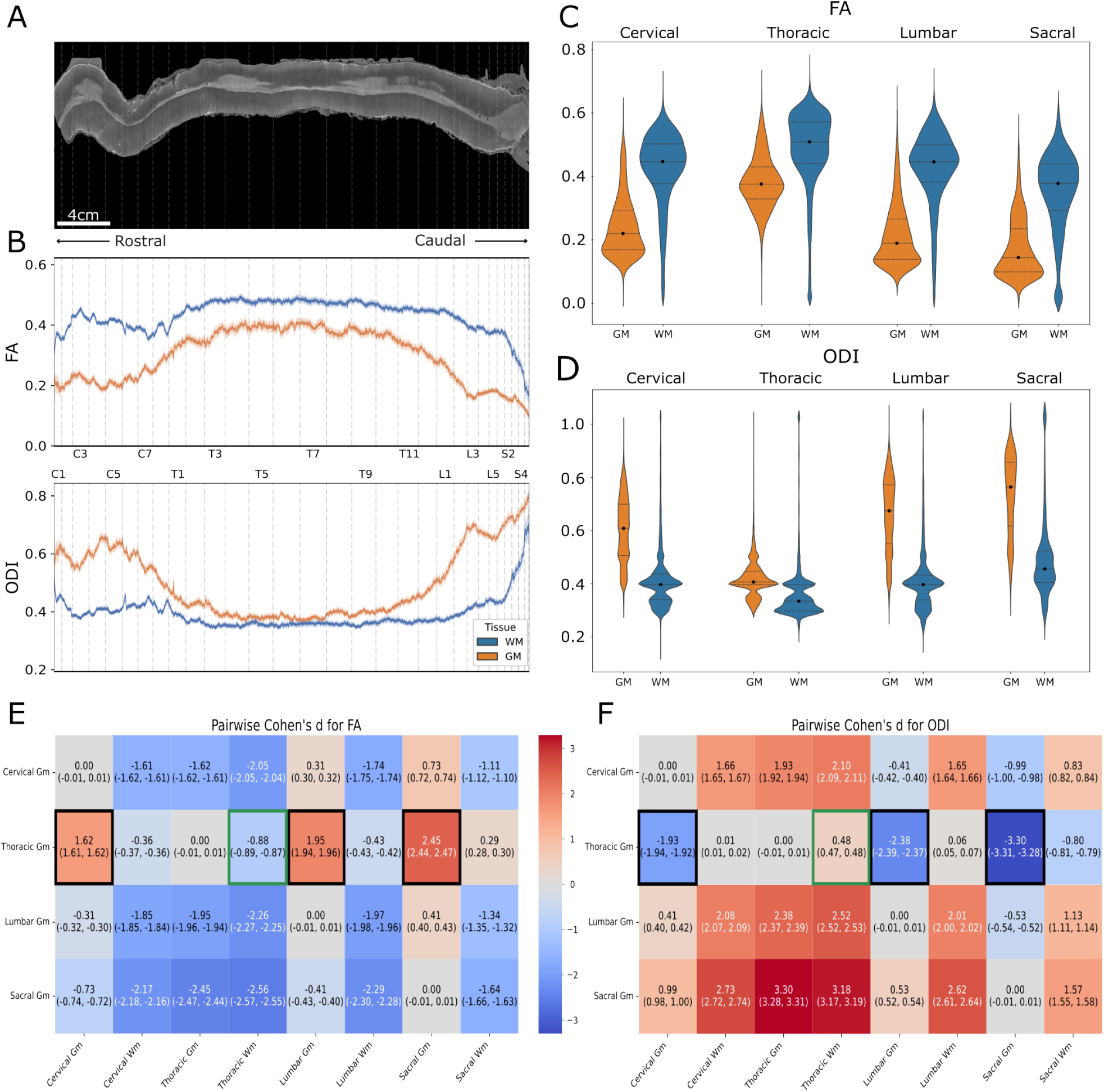
**Fig A.** T2 sagittal slice, stippled white lines demarcates segment boundaries. **Fig B.** Rostral-Caudal gradients of top panel; FA and lower panel; ODI. **Fig C & D**. Probability density distribution of FA (**C**) and ODI (**D**) for the four spinal segments and tissue types. Lines denote the median together with 25% and 75% quantiles. **Fig E & F.** Effect size heatmaps of (Cohen’s d) summarizing pairwise differences across tissue types and spinal segments. Key comparisons are highlighted with a black (GM) or green (WM) ROI box

#### Constrained spherical deconvolution reveals a dominant rostro-caudal orientation in thoracic gray matter

DTI and NODDI have provided scalar measures of anisotropy and dispersion but are incapable of elucidating the full scope of the organization of fiber directionality across the hemisphere. We therefore applied constrained spherical deconvolution (CSD) to estimate the full fiber orientation distribution function (fODF) in each voxel (Fig. 3). The multi-tissue RUMBA-SD algorithm (38) was used to mitigate partial-volume effects at the GM–WM interface, with separate response functions estimated for WM, GM, and the perfluorocarbon fixation buffer. The fODF glyphs visualized across representative axial sections of all four major spinal segments reveal a striking contrast between thoracic gray matter and the other regions (Fig. 3A). In cervical, lumbar, and sacral gray matter, the glyphs are predominantly green and red, indicating lateral and dorso-ventral orientations dominating within the transverse plane, a pattern consistent with the known radial dendritic arborizations and commissural projections of spinal interneurons. The thoracic gray matter, by contrast, is overwhelmingly populated by blue glyphs, indicating that the principal fODF lobes point in the rostro-caudal direction with no transverse components showing as their relative size to that of the rostral-caudal direction being minuscule. Close inspection of the commisural line does depict few minor lateral glyphs. Voxels with a multi-orientation glyphs representation bordering white matter and the perfluorocarbon fixation buffer at the most ventro-medial part of the thoracic slices were deemed to be either an artefact or a ventral root nerve. We therefore excluded these voxels from further analysis as they were not thought to be representative of neither WM or GM. The visual observation of the gray matter segment specific variation in orientation distribution is confirmed quantitatively in the azimuth and polar angle rose plots (Fig. 3B). All diffusion orientations **v** are defined on the unit sphere S^2^ ⊂ R^3^. however, since diffusion is symmetric that is, diffusion along **v** is indistinguishable from diffusion along −**v** the effective domain reduces to the projective plane RP^2^, which is equivalently represented as the upper hemisphere S^2^. To represent something three dimensional in a two-dimensional world we parametrized orientations in spherical coordinates (θ, ϕ). However, the azimuth angle ϕ is degenerate at the poles (θ = 0^◦^, 180^◦^), where the transverse component vanishes and ϕ becomes undefined. Analyses relying on ϕ were therefore restricted to orientations satisfying θ > ɛ for a small threshold ɛ = 85^◦^. The thoracic gray matter θ is highly centered around 90°, with 60.4% of the orientations falling in between 85°-95° θ. With an additional 16.2% in between 80°-85°, and only 23.4% with an θ >10°. The polar angle distribution of the remaining segments all exhibit significantly fewer orientation centered around the pole. With cervical GM having the second most, and sacral GM exhibiting the fewest number of rostral-caudal orientations. We further analyze the gray matter ϕ distributions for the orientations with an θ > ɛ across spinal segments. The binarized distribution of θ of can be seen in (Fig. 3C). The binarized fODF vector ratio (Fig. 3C) provides a direct, segment-level summary: 91.7% of cervical fODF vectors are transverse in orientation, as are 95.6% and 97.1% in lumbar and sacral segments, respectively. In thoracic gray matter, this proportion drops to 39.6%, with the remaining 60.4% classified as near-pole, i.e., oriented strictly toward the rostro-caudal axis. We observe a thoracic azimuth distribution notably more uniform and less structured than those of remaining segments. With thoracic GM exhibiting two principal azimuth orientations, (1) lateral, (2) dorsal-ventral. Cervical, lumbar and sacral gray matter are on the other hand all composed of a more spread out azimuth orientation distribution. The full analysis of WM and GM alike along with a flattend hemispheric heatmap representation of the full fDOF can be found in S. 13-16. These findings demonstrate that a substantial population of neurites in the thoracic gray matter runs longitudinally along the cord axis, a configuration that departs markedly from the classical view of gray matter as a region of transversely organized local circuity we observe at cervical, lumbar and sacral regions. We further aim to elucidate and verify our findings using fluorescence microscopy and tagging non-phosphorylated neurofilament H proteins using SMI32. Our microscopy analysis is limited to the 2D transverse plane and we would thus expect to see limited transverse projection or very short transverse projection for a pixel-wise structure tensor analysis of the thoracic gray matter, with more prominent in plane projections for the remaining gray matter segments.

**Figure 3:**
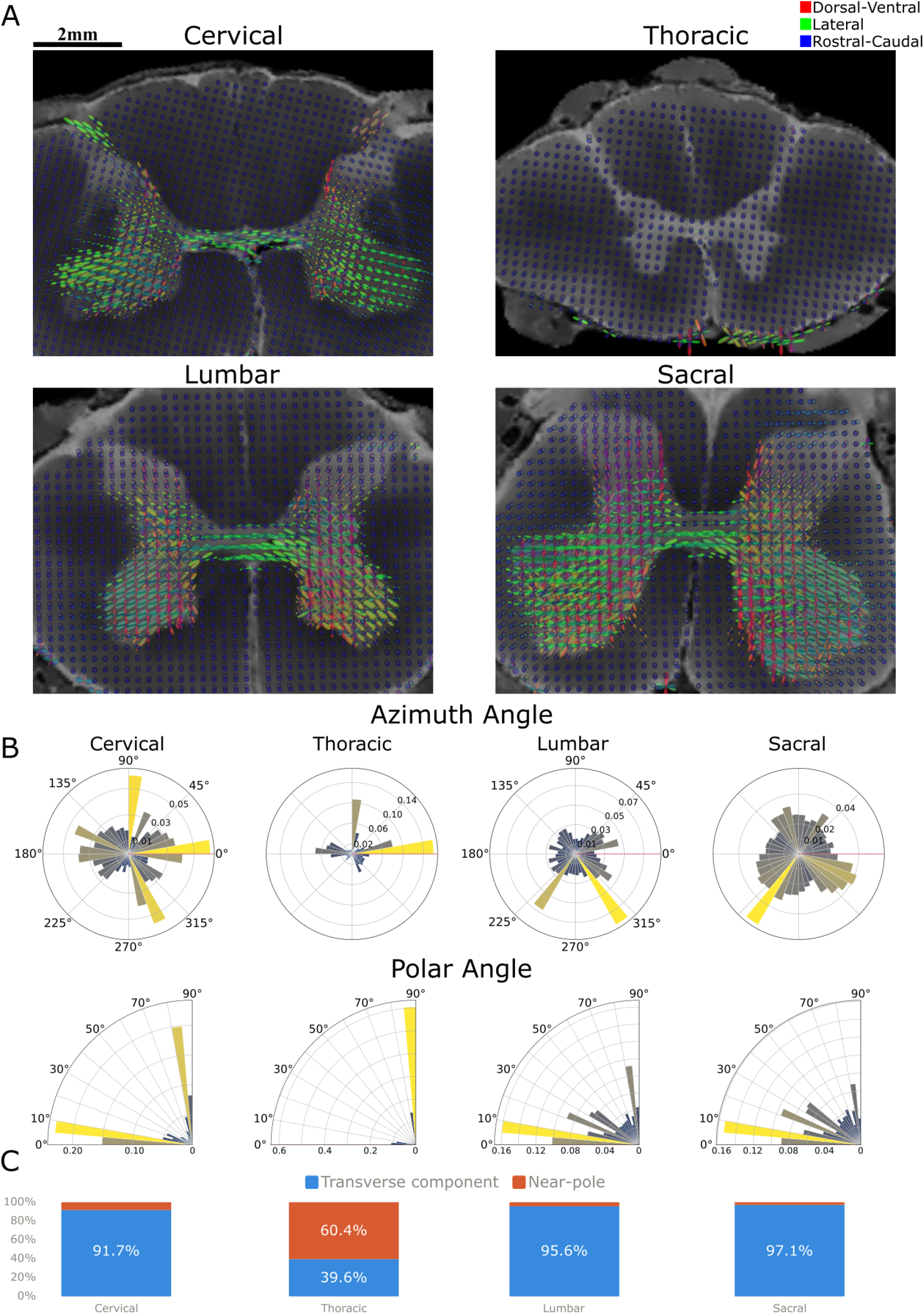
**Fig A.** Fiber orientation distribution per voxel depicted by 3D rendered glyphs. Orientation between the cartesian directions are blended. **Fig B.** Distribution of gray matter fODF orientations on an hemisphere. **Fig C.** binarized ratio of gray matter fODF vectors with a polar angle >85 and < 95 into longitudinal fODF vs transverse component

### 2.2 Microscopy

We collected fluorescence microscopy images to investigate the neurite microstructure at an resolution exceeding that of most axonal widths, in addition to serving as a validation of the rostral-caudal fODF profiles we exempted in the previous section. After removal of the dura mater, the spinal cord was divided into its four regions and cut into 50 µm transverse sections on a microtome. Free-floating sections underwent immunohistochemistry for non-phosphorylated neurofilament H (SMI-32), together with NeuN, GFAP, S100 and ChAT, and were counterstained with Hoechst; full antibody and staining details are given in the supplementary methods. Sections were imaged on a Zeiss Cell Observer spinning-disc microscope in widefield mode using a 20× 0.8-NA objective, with SMI-32 visualized via an Alexa-Fluor 647 secondary. Approximately 60–80 z-stacks were acquired per segment. Due to uneven sectioning, large portions of these volumes were out of focus causing degrading contrast, SNR and preventing robust phase-correlation stitching. Consequently only 8–10 slices per z-stack retained more than 80% of the axial plane in focus. Robust structure tensor reconstruction of longitudinal projections was therefore not possible, and the microscopy analysis was restricted to the 2D transverse plane. To address detector noise, photon shot noise and background noise we deployed Noise2Void (N2V), a self-supervised deep-learning denoising framework (39), producing state of the art denoising directly from single noisy images without the need of a clean version. We trained a version of N2V for each of the images using a patch size = 512×512 and a ROI of size 8. Local fibre orientation and structural anisotropy were estimated from fluorescence images using a 2D structure tensor framework. We tested multiple combination of structure tensors parameters but ended up using a structure tensor **S** parameterized towards find grained pixel level orientation flows using a gradient smoothing scale σ= 2.5 px and an integration scale ρ = 10 px. The technical details regarding structure tensor and our choice of paramters can be found in the supplementary material under 6.5.

**Figure 4:**
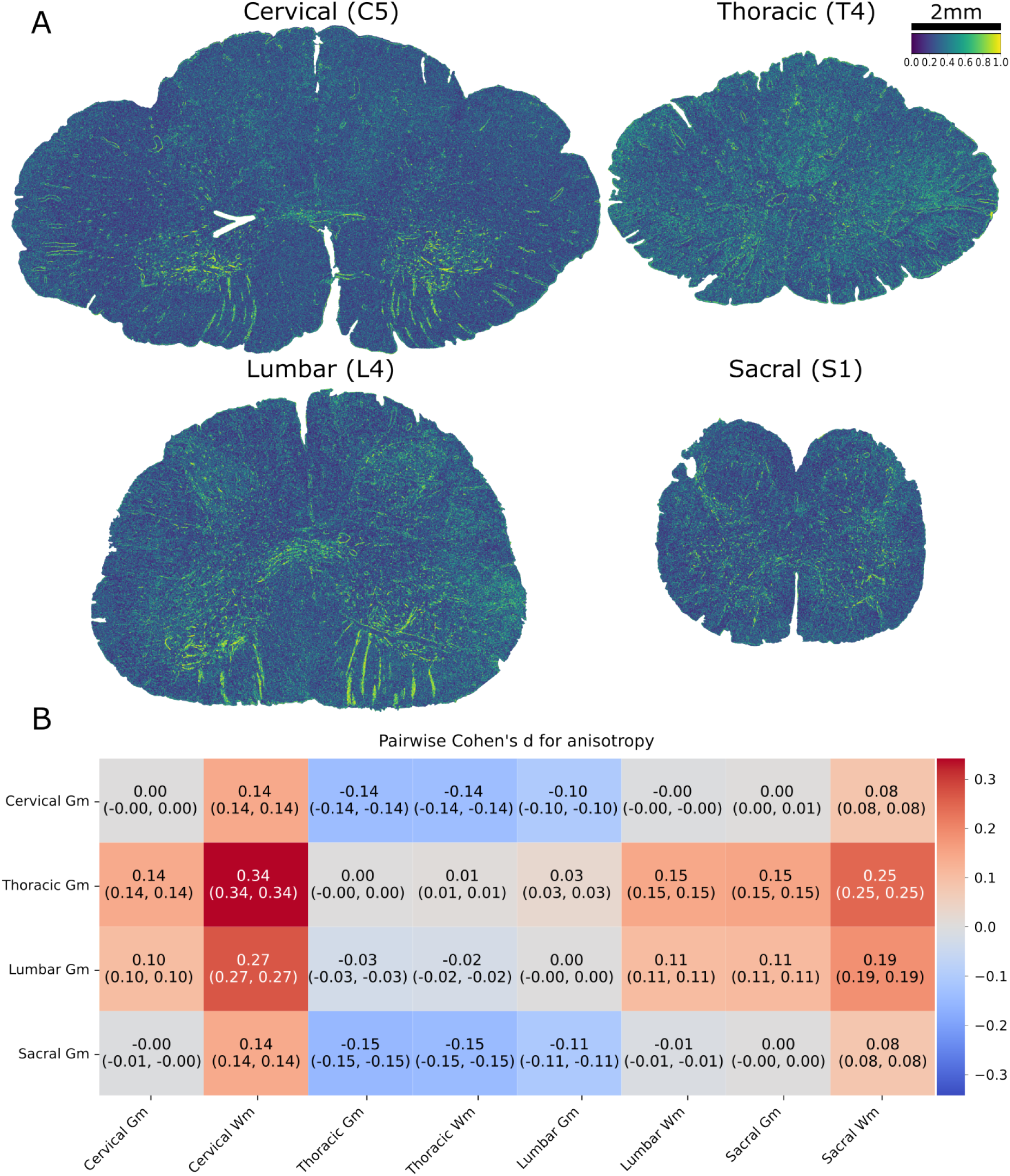
**Fig A.** Anisotropy dervied from the structure tensor model of the of the fluorescent microscopy SMI32/AF647. **Fig B.** Pairwise Cohen’s D effect size heatmap of ST anisotropy grouped by tissue + segment

#### Pixel-wise structure tensor anisotropy maps

Methodological differences between structure tensor and dwMRI entails that the anisotropy derived from our structure tensor is not a 1:1 match of that observed from our dwMRI. The signal source differs fundamentally. FA/ODI are *indirect* proxies inferred from the displacement profile of water, the structure tensor measures orientation directly from the spatial gradients of the labelled structure itself. So one is a diffusion-mediated estimate of geometry, the other a measurement of the geometry as imaged. Furthermore, FA/ODI integrate over the full 3D volume of a 200×200×200 µm voxel, capturing through-plane orientation. Whereas structure tensor anisotropy is computed in-plane over a 2D neighbourhood orders of magnitude smaller. A fibre population that is coherent in 3D but obliquely sectioned, or coherent in-plane but dispersed through-plane, will be scored differently by the two methods. While a direct comparrison between the two are meaningless, they do still carry complementary information in regards to the neurite microsctructure (40,41). With microscopy resolving the actual in-plane fiber arrangement at sub-micron resolution.

Across all four segments 4a, high-anisotropy values were concentrated along coherent linear structures, most prominently the ventral and lateral white matter fascicles and ventral root fibers, which appear as bright green-to-yellow streaks against a low-anisotropy (∼blue) parenchymal background. The gray matter portion was comparatively low in anisotropy for cervical, lumbar and sacral sections, consistent with the multi-directional, radially arborising local circuitry expected of these segments. The thoracic gray matter departed from this pattern with in-plane anisotropy being the highest of the four gray matter segments. This might seem counter intuitive since their orientation is longitudinal. But reduced variance in the intensity gradients directions arises as a consequence of strictly coherent fiber bundles, also in plane. Whereas an in plane overlapping with fibers with orientations twisting and turning causes reduced anisotropy, longitudinally oriented neurites sectioned transversely can produce coherent, elongated cross-sectional profiles. To quantify the segment- and tissue-level differences visible in the pixel-wise maps, we summarized segment wise mean anisotropy with pairwise effect sizes (Cohen’s d), using the same framework applied to the dwMRI metrics. As before, classical null-hypothesis tests were uninformative given the large number of data points per group, so d was used as a direct measure of effect magnitude, with 95% confidence intervals from the large-sample approximation and effect sizes interpreted against the conventional thresholds (small |d| < 0.5, medium 0.5 ≤ |d| < 0.8, large |d| ≥ 0.8). The resulting effect sizes are shown as a diverging heatmap in 4b. The defining feature of this comparison is the magnitude of the effects: every pairwise difference in in-plane anisotropy fell within the *small* range (|d| < 0.5), with the largest effect reaching only d ≈ 0.34 in thoracic gray matter. Thus showing similair trends to the FA/ODI analysis but with clear differences in terms of the effect size scale. In similar fashion to the analysis of dwMRI based metrics we also conducted an in-depth analysis of the spatial autocorrelation using Moran’s I and its effect on our dependent variable in addition multiple-comparison correction. The analysis verified our observations and can be found in S. 17. As mentioned, crucially the dwMRI and fluoresence microscopy investigate at two different spatial scales and biophysical phenomenons, a 1:1 match in effect sizes are therefore not expected. Taken together, the structure tensor results corroborate the dwMRI findings on independent, directly labeled SMI-32^+^ neurofilament and at micron resolution. The comparatively dramatic thoracic distinctiveness seen in 3D (FA, ODI, and the rostro-caudal fODF) collapses to a small in-plane effect when the measurement is restricted to the transverse plane.

#### Anisotropy-weighted orientation maps

To resolve the direction of in-plane organisation, we mapped the pixel-wise structure tensor orientation θ = arctan2(v*_x_*, v*_y_*) onto a cyclic colour wheel and modulated colour intensity by the local anisotropy via the alpha levels. Coherent high-anisotropy fibres render as vivid, hue-coded streaks and low-anisotropy regions are dimmed 5. A region of gray matter was selected per segment and shown at high magnification. The four segments separate into two qualitatively distinct regimes. Cervical, lumbar and sacral gray matter were populated by abundant elongated, coherently oriented fibres spanning a range of in-plane angles, visible as bright, multi-colored i.e., multi directionaltion continuous streaks. Lumbar and sacral gray matter in particular showed dense oblique fiber bundles spanning multiple orientations, constituting transverse local circuitry. Thoracic gray matter, by contrast, was strikingly devoid of coherent linear structure, instead it was dominated by a fine, uniform punctate texture with only sparse elongated fibres 5A. This punctate pattern is the expected in-plane signature of fibres sectioned transversely, i.e. of a neurite population running out of the imaging plane along the rostro-caudal axis, and stands in direct contrast to the elongated, in-plane fibre streaks seen in all other segments. The anisotropy-weighted orientation distributions are summarised as polar histograms for gray and white matter across the four segments (Fig. 5B). Whereas 5A provides information about the orientations of the individual neurites, 5B instead asks what the orientation distribution looks like for gray and white matter per segment. While images in Fig. 5 are cropped to regions of interest the orientation distributions in 5B are a depiction of the entire uncropped cross-sectional slice. Distributions profiles ranged from strongly bi-lobed polar distributions indicating a well-defined preferred in-plane axis, to near circular distributions, indicating an absence of any dominant in-plane orientation across the segment. Strikingly, the thoracic gray matter distribution was similar across gray and white matter, both with strongly bi-lobed polar distributions, around a single preferred in-plane axis. Although the thoracic neurites are sectioned transversely and therefore appear punctate rather than streak-like, their asymmetric cross-sectional profiles share a highly consistent in-plane orientation due all longitduinal axons being cut at a similar angle, yielding a more coherent orientation distribution than the in-plane fiber populations of lumbar and sacral segments. Whom by contrast showed in-plane fibers span multiple crossing orientations in 5A, producing comparatively broader, less concentrated distributions. The polar profile for cervical gm is note worthy because it shows a similar distribution to that of thoracic gm, but instead its bi-lobed profile is centered around 0*, constituting lateral orientations. Whereas the bi-lobed thoracic profile arose due to short non-linear neurite patterns the cervical gm distribution arises as a consequence of the proportional increase in size of the commissarial midline, whose projection profile is pre-dominantly literalized.

**Figure 5:**
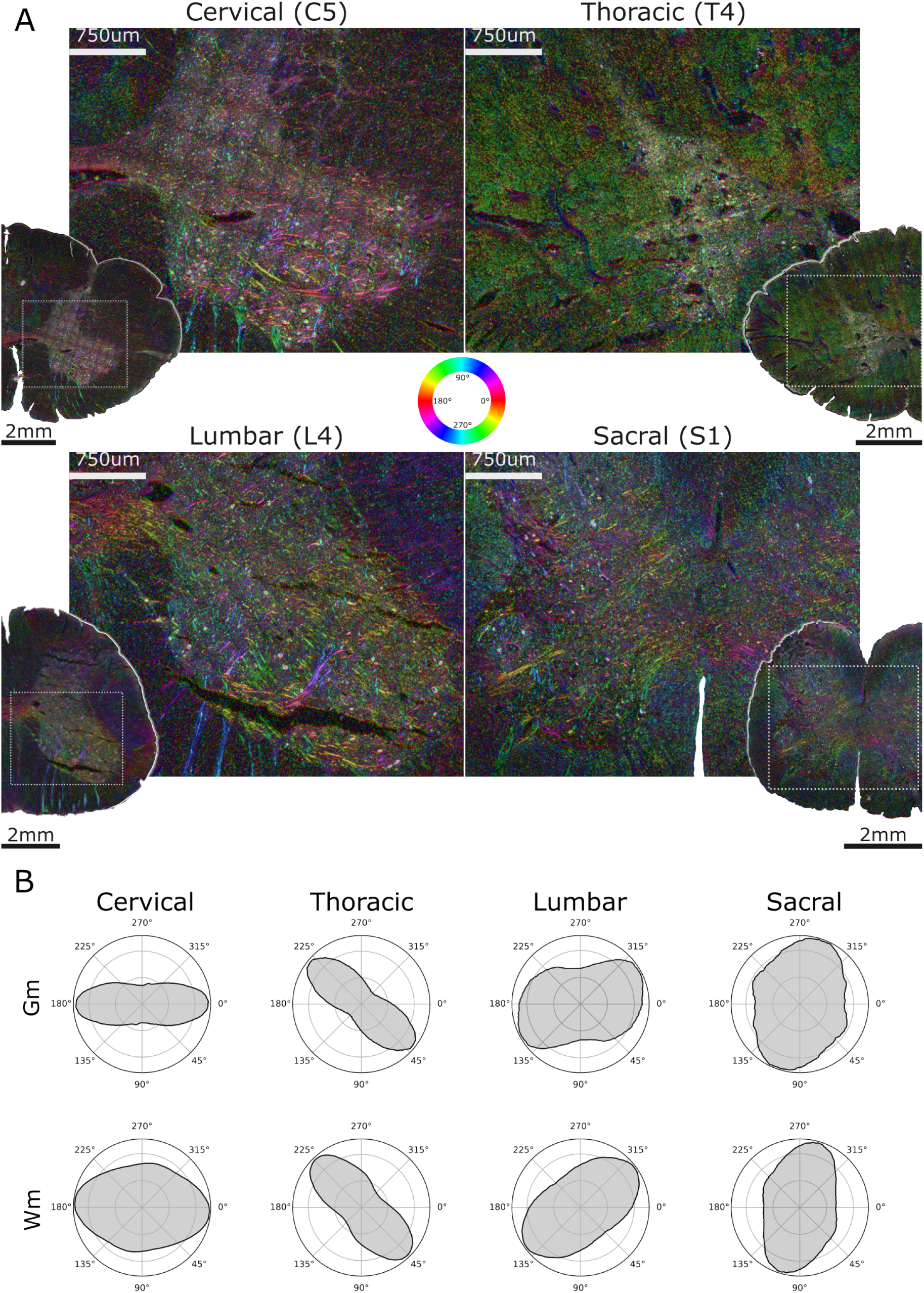
**Fig A.** Fluoresence microscopy images of SMI32/AF647 overlayed with the anisotropy weighted orientation color scheme shown on the color wheel. **Fig B.** Polar plots of the circular orientation statistics for gray matter and white matter.

#### Structure tensor orientations and dwMRI fODF converge on a shared regional orientation architecture

As a final, cross-modal validation we illustrate similairites with a side by side view of the dwMRI fibre orientation distribution functions and the structure tensor orientations from fluorescence microscopy. With the left panel depicting fODF glyph field overlaid on the structural image (left) and to the right the anisotropy-weighted structure tensor orientation map (right; Fig. 6). The two modalities are not numerically equivalent, the fODF is an indirect, model-based estimate integrated over the full 3D diffusion profile of a 200 µm voxel, whereas the structure tensor directly measures in-plane gradient orientation in a 2D section at 0.325 µm/pixel, so the comparison tests agreement on the dominant orientation of each region rather than a one-to-one correspondence of values. In the fODF colour scheme, blue denotes rostro-caudal, green lateral, and red dorso-ventral orientations, with intermediate hues (yellow, magenta, cyan) encoding the corresponding oblique combinations. In cervical, lumbar and sacral gray matter the two modalities agreed. The fODF contained substantial in-plane structure green (lateral), red (dorso-ventral) and oblique magenta/cyan glyphs, including the prominent ventro-lateral fan of lateral glyphs corresponding to the ventral white matter and root fibres and the structure tensor maps reproduced these in-plane orientations as coherent, matching fibre streaks. Where the dwMRI reported fibres lying within the transverse plane, the microscopy recovered them concordantly.

**Figure 6:**
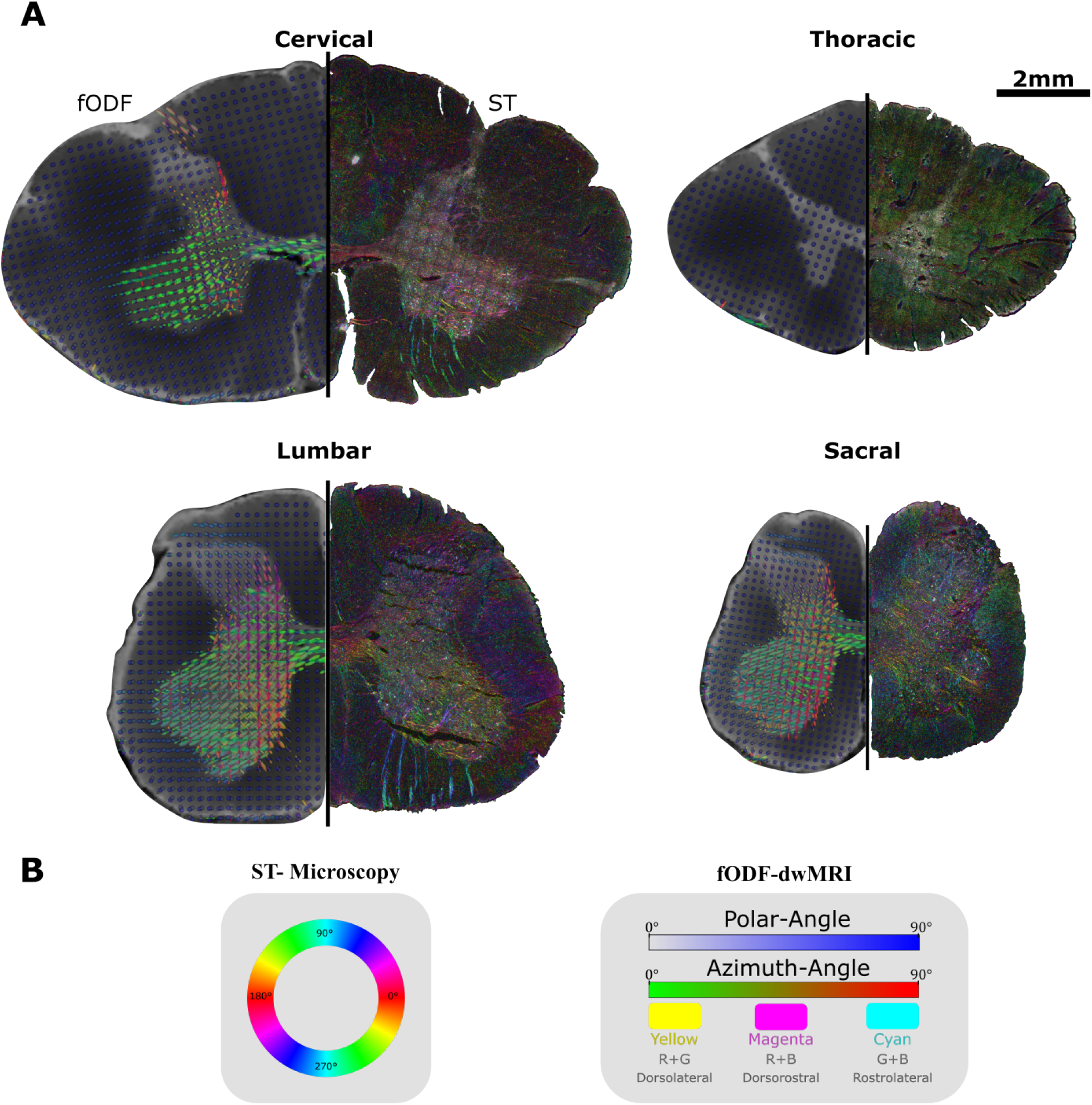
Side by side visual comparison of fODF from MRI and structure tensor orientations from the fluorescence microscopy. Associated color schemes are depicted below with (left) orientation wheel mapping onto ST-microscopy orientations and (right) color bars mapping to the fODF glyphs orientations.

The thoracic gray matter is the decisive case, and here the two modalities are consistent in the only way their dimensional difference permits. Its fODF was overwhelmingly dominated by blue glyphs rendered as small dots because the lobes point out of the imaging plane indicating a near-pure rostro-caudal orientation across almost the entire gray matter, mirroring the binarised estimate that 60.4% of thoracic orientations fall within 85–95^◦^ of the rostro-caudal axis (Fig. 3. The matched structure tensor map showed no corresponding long coherent in-plane neurites, instead it displays the same fine punctate, high-anisotropy texture seen in Fig. 5, i.e. the transverse cross-sections of those longitudinal fibres. The agreement is therefore one of mutual constraint, the dwMRI specifies the rostro-caudal axis that the 2D microscopy cannot directly render, and the microscopy confirms, at the level of directly labelled SMI-32^+^ neurofilament, that essentially no in-plane fibre structure remains to account for the thoracic signal otherwise. Taken together, Figs. 5 and 6 establish that the regional orientation architecture inferred from ex vivo dwMRI is reproduced by an independent histological modality with orthogonal assumptions. Both converge on the central finding that thoracic gray matter is distinguished from every other spinal segment by a predominantly rostro-caudal neurite orientation, while the remaining segments are characterised by transversely organised, in-plane fibre populations.

## 3 Discussion

In this study, we used two complementary methods to establish the fiber orientation of primarily human spinal cord gray matter. The main observation of both methods is that the fiber orientation of the thoracic gray matter is radically different from the other spinal regions because it is predominantly in the longitudinal directions. In contrast the gray matter within the lumbosacral and cervical regions is predominantly in the transverse direction. We regard three non-exclusive substrates as the most plausible candidates for the observed thoracic gray matter rostro-caudal neurite orientation in contrast to the radial patterns observed in the remainder of the cord, and we emphasise that the present data constrain geometry but not cellular identity, which would constitute a new study in itself. First, the segmental range of the effect coincides with the intermediolateral cell column (IML), which in humans is confined to the thoracolumbar cord, approximately T1-L1/L2. Sympathetic preganglionic neurons of the IML are classically described as forming rostro-caudally arranged clusters whose dendrites are oriented predominantly along the long axis of the cord, producing extensive longitudinal dendritic bundles and a ladder-like distribution about the central canal (42–44). This morphology is close to an anatomical definition of what we measure as a low-dispersion, pole-directed orientation distribution. That our axial-diffusivity and FA maxima fall around T5-T8, near the middle of the IML’s extent rather than at its borders, is consistent with a contribution that scales with the density of this architecture. Second, Clarke’s column (nucleus dorsalis) spans approximately C8-L3 and is a longitudinally continuous nucleus giving rise to the dorsal spinocerebellar tract. Golgi studies report that the dendritic arborisations of its principal cell types extend mainly in the longitudinal direction, remaining within the nucleus and spreading up to a millimetre (45). We note that this picture is however not uncontested. With other classification of Clarke’s neurons in the adult cat described as multipolar cells with radially projecting dendrites and fusiform cells whose long axis is perpendicularly oriented (46). So the combined net orientation contribution of this nucleus is likely to be mixed rather than purely axial. Additionally would laminar or nuclei specific rostro-caudal projecting dendrites from Clarke’s nucleus or sympathetic preganglionic neurons of the IML not explain the gray matter wide longitudinal orientation profile we observe. As their contritubtion would be far too local to drive the macro scale phenomenon we observe across all lamina. Third, and more prosaically, the thoracic cord has the smallest gray-matter cross-section of the cord (47) and the lowest density of motor neurons and limb-related pre-motor circuitry of any region (48). Long propriospinal and descending fibres that traverse or skirt the gray matter are therefore diluted by comparatively little transverse local arborisation. On this account the thoracic signature might be less the addition of a longitudinal population than the subtraction of the transverse one. These explanations differ in their predictions, with IML or Clarke’s contribution predicts a spatially clustered, laminar-specific effect, whereas dilution predicts a functionally spatially uniform one across the entire thoracic gray matter. With the latter conforming better to our observed data. As previously motivated, if projection geometry is taken as a proxy for the routes along which spinal circuits exchange information, the regional dissociation we observe maps onto a functional dissociation that is independently well motivated. The cervical and lumbosacral enlargements serve distal, high-degree-of-freedom limb musculature, and are the segments in which encephalon-independent rhythm and pattern generation has been most convincingly demonstrated (13,18). A transversely wired, high-dispersion gray matter is what one expects of tissue performing dense, segment-local computation. Exactly what we observe in our data. The thoracic cord, by contrast, serves axial and postural musculature and the sympathetic outflow, both of which plausibly require coordination *along* the neuraxis rather than within a segment graded sequential recruitment of trunk muscles, and the rostro-caudally organized topography of sympathetic control. A longitudinally dominated architecture is therefore the system geometry appropriate to that task. A bulk of plausible counterfactual explanations for our observation are inevitable and it should be the focus of a future study to more formally test the causal factor for the observations made in the this study.

## 4 Conclusion

We set out to ask whether the organization of neuronal projections in the human spinal cord varies systematically along its rostro-caudal extent, beyond the classical demarcation of the longitudinal white-matter tracts. Combining ex vivo high-angular-resolution dwMRI across segments C1-S5 with structure-tensor analysis of SMI-32/AF647 fluorescence microscopy, we find that it does, and that the variation is continuous and gradual along the cords length with categorical segment level characteristics. Cervical, lumbar and sacral gray matter with low fractional anisotropy, high orientation dispersion, and fiber orientation distributions dominated by transverse components (91.7%, 95.6% and 97.1% of fODF vectors respectively), reproduced at micron resolution as dense, multi-directional in-plane neurite streaks. Thoracic gray matter does not. It shows elevated axial diffusivity peaking near T6, an FA distribution shifted upward (x̃ = 0.35) into the range otherwise occupied by white matter, an ODI distribution both narrower and lower (x̃ = 0.39) than any other gray matter region, and an fODF in which 60.4% of orientations fall within 85-95^◦^ of the rostro-caudal axis. The microscopy is concordant in the only way a transverse section can be: thoracic gray matter is devoid of the long, coherent in-plane neurites seen elsewhere, and is instead dominated by a short punctate texture, the expected cross-sectional signature of a fiber population running out of the imaging plane. The central discovery of this work is therefore that thoracic gray matter is distinguished from every other gray matter region of the human spinal cord with previously unreported predominantly longitudinal neurite architecture, in contrast to the highly transverse lumbosacral and cervical gm, and that this is visibly and statistically verified with two modalities. We further hypothesize about the advantages and neuro-anatomical explanations for our observed changes in neurite microstructure, surmising functional-anatomical coupling between the spinal segments functional tasks and the requirements of the systems architecture best equipped to facilitate the given function.

## Code and Data availability

A clean version of the code is available on GitHub, for all scripts including experimental tests contact the main author. MRI data is available at DOI, and fluorescence microscopy data is available upon request.

## Author contributions

- Sigurd Fyhn Sørensen: Writing the main and supplementary manuscript along with preprocessing, data modeling and analysis.
- Jaspreet Kaur: Collection and data preparation for fluorescence microscopy data.
- Rune Berg: Supervision and conceptualization of the study.

## 5 Supplementary Methods

In this section we provided a detailed outline of everything from preprocessing and modeling choices to statistical tests and additional analytical methods. We divide the section into three parts, part one deals with MRI and associated data acquisition, preprocessing and modeling, part two contains similair elements to part one but with a focus on the microscopy. Finally the last section dives into the statistical tests used in the paper across both data modalities.

## Magnetic Resonance Imaging

### 5.1 Data Collection

This study was performed on a full-length skull stripped spinal cord from a deceased individual who had bequeathed her body to science and education at the Department of Cellular and Molecular Medicine (ICMM) of Copenhagen University according to Danish legislation (Health Law No. 546, Section 188). Furthermore, the study was approved by the head of the Body Donation Program at ICMM, ensuring ethical approval. The spinal cord originated from a 91-year-old Caucasian female without any known neurological or psychiatric diseases. It was obtained, dissected, and fixed within 24 hours post-mortem. In order to preserve tissue morphology, an immersion fixation protocol was performed using a paraformaldehyde (4%) buffer with a pH of 7.4 (26, 27). The tissue was fixed in the buffer for 2 weeks at low temperatures. For long-term storage, it was transferred to a phosphate-buffered saline solution with a pH value of 7.4. Images were acquired on a 9.4 T preclinical MRI system (BioSpec 94/30; Bruker Biospin, Ettlingen, Germany) equipped with a 1.5 T/m gradient coil. Prior to imaging, the spinal cord was placed in a plexiglas tube and immersed in fluorinert (FC-40, Sigma-Aldrich), reducing background signal Our protocol involed immersing the ex vivo tissue in perfluorocarbon prior to scanning. As perfluorocarbons contain no hydrogen protons they produce negligible background signal for conventional hydrogen-1 MRI. Producing enhanced contrast between tissue and background, increasing SNR and CNR (28). Perfluorocarbons furthermore have magnetic susceptibility close to that of tissue, reducing susceptibility artifacts at tissue boundaries, in adition to being a dense and inert fluid therby providing physical stabilization and mitigation of tissue heating (29, 30). A limited field of view (1.6 cm) relative to the length of the spinal cord (40 cm), necessitated imaging being done in 29 sections, known as Multiple Overlapping Thin Slab Acquisition (MOTSA) MRI (31, 32). Between each section-scan the spinal cord was advanced 1.4 cm by a custom-built mechanical stepper, resulting in a 0.2-cm overlap between neighboring sections. Structural MRI were acquired for each section with a T2-weighted 2D RARE sequence with associated scanning parameters, echo time (TE) = 30ms, repetition time (TR) = 7000ms, a field of view of 1.92 × 1.92 × 1.6 cm^3^, and a matrix size of 384 × 384 × 80, resulting in 50 × 50 µm^2^ in-plane resolution and a slice thickness of 200 µm, producing an anisotropic voxel of size 5e5 µm^3^. The anatomical T2 images were generated using 20 excitations, i.e., traversing k-space 20 times. Thus,the final image is an average of the 20 excitations. Doubling the number of excitation (NEX) doubles the scan time but also improves SNR by a factor of √2. Increasing NEX and the number of averages is therefore the most fundamental way to increase SNR if the study permits prolonged scanning time. A NEX of 20 is incredibly high with a total scan time of 400 hours, and only possible due to the fact that our study is conducted on ex vivo soft tissue. The dwMRI was also acquired for each of the 29 sections with a spin-echo DWI sequence. We used a single shell with a b-value = 4000s/mm^2^, along with eighty motion-probing gradients, encoding 80 different diffusion directions. Yielding eighty b-vectors on a hemisphere, along with three b0 images corresponding to a T2-weighted sequence. The three b0 images were averaged to create a single optimal b0 image. With eighty diffusion directions our data constitute what is known as high angular resolution diffusion images (HARDI) (49–51). Allowing for demarcation of small-angle changes in micro-structural diffusion. Scan parameters were as follows: echo time (TE) = 17.2ms, repetition time (TR) = 5000ms, a field of view of 1.92 × 1.92 × 1.6 cm3, a matrix size of 96 x 96 x 80, producing isotropic voxels with 200 x 200 µm m2 in-plane resolution and a slice thickness of 200 µm, giving us a voxel size of 8e6 µm^3^. The spin-echo DWI sequence used a NEX of 1 as we expected the perfluorocarbon immersion to be sufficient in facilitating a workable SNR. All sequences used single-phase encoding direction (A-P) for all of the sections. Dual-direction phase encoding can be advantageous for DWI/DTI, particularly when using EPI sequences in live subjects. Its’ effect on image quality and accuracy however is typically minor in postmortem scans as fixed samples exhibit negligible motion or flow. To facilitate future image stitching between sections, we intentionally chose a spin-echo DWI sequence with a short echo time (52). This choice minimizes susceptibility to eddy current artifacts, which can be more prominent in sequences with longer echo times (53). Making co-registration between adjourning sections more stable. Finally the scanner’s gradient power of 1500mT/m, allowed the flexibility to design optimized pulse sequences adjusting the rise time and duty cycle towards minimizing eddy currents related to magnetic switching (54). The 29 slabs of dwMRI and T2 were exported as DICOM files from the scanner software (ParaVision 360, Bruker) and converted to NIfTI format using open-source software dcm2niix, (55).

### 5.2 Preprocessing

MRI data is inherently noisy, especially at higher field strengths, requiring several preprocessing steps in order to extract useful quantitative analytical information (56, 57). As reported by (58) choices in the preprocessing pipeline have direct cascading effects on analysis results. Intra-lab variability in preprocessing pipeline therefore has potential affects in terms of reproducibility. This is further complicated by the fact that no single preprocessing pipeline being optimal for all datasets. The following section, therefore, aims to provide a detailed description and justification for our choice of preprocessing, with sufficient detail to facilitate easy replication. Our choises were weighted based on two principles, (1) a too-small preprocessing pipeline can fit noise as an actual signal, (2) an over-processed image introduces artficial bias and artifacts causing alterion of image features and an inherent reduction in reproducibility of subsequent analysis. We therefore aimed at producing preprocessing steps that improved signal quality while minimizing artificial bias. All preprocessing steps were independently applied to each slab prior to stitching for the DW-MRI and T2 data.

#### 5.2.1 Denoising

We utilized an Marchenko–Pastur PCA (MP-PCA) algorithm for anatomical T2 denoising. The method requires high redundancy in the input data to effectively separate the signal (low rank) from the noise (high rank). Making it ideal for our dwMRI data with signal across multiple volumes, diffusion directions and echoes in a low-rank subspace. Suppressing thermal and physiological noise while preserving microstructural details. The algorithm has been shown to provide an optimal compromise between noise suppression and loss of anatomical information for various techniques, including DTI (Manjón et al., 2013), spherical deconvolution (Veraart et al., 2016), and DKI (59). We utilized an algorithmic implementation in DIPY (60), a Python library for diffusion dwMRI, with a patch-radius parameter of two around each voxel. Resulting in patches with a size of 5×5×5. T2 images were denoised using a non-local means denoising algorithm (nlmeans) developed by (61). We ran the Nlmeans denoising with a patch radius of 2 voxels and a block radius of 5 voxels, with an assumption of the noise being Rician noise.

#### 5.2.2 Eddy Currents

Eddy currents are caused by rapid gradient switching, leading to possible nonlinear geometric distortions of dMRI images. Said distortions vary with gradient direction and strength, thereby slightly misaligning the different diffusion volumes. Voxel-wise analysis, e.g., tensor fitting, tractography, and orientation distribution functions (ODF), require DW-MRI images to be mapped on the same anatomical space (62). Individual diffusion scans are therefore co-registered to the averaged b = 0 image using advanced normalization tools (ANTS) (63). Correcting for B0 drift and eddy-current distortions. Utilizing dynamic field correction, more precisely an affine registration with shear and scale terms along with 12 degrees of freedom and an entropy-based metric to deal with contrast differences between the reference B0 and diffusion gradient images.

#### 5.2.3 Gibbs Correction

Gibbs Correction: Gibbs artifacts, also known as truncation, ringing, or spectral leakage artifacts, can appear as multiple fine parallel lines immediately adjacent to high-contrast regions. For example, in the boundary between white-matter and gray matter or background. Spinal imaging has proven to be problematic due to these artifacts, as they may artifactually widen or narrow the cord or mimic a syrinx (64). Gibbs artifacts arise due to the use of Fourier transforms used to transform the raw MR signal into images. Theoretically can any signal be accurately represented by an infinite summation of sine waves with different amplitudes, phases, and frequencies. However a restricted sampling to a finite number of frequencies in MR imaging leads to an approximation of the image using relatively few harmonics of its complete Fourier representation. The finite Fourier series therefore truncate the full information leading to loss of infomation (65). Standard filtering approaches used to overcome this inevitably also reduce the effective image resolution. We therefore implemented the Gibbs correction algorithm from (66) using default model parameters. The technique is based on local sub voxel-shift resampling. Rather than applying the traditional spatial smoothing, leading to a reduction in image sharpness, the algorithm locally estimates the voxel shift that minimizes total variation within a small neighborhood. Thereby only introducing bare minimum alteration of the image needed. The image is then resampled at these optimized positions, effectively suppressing Gibbs oscillations while preserving anatomical edges and maintaining effective resolution.

#### 5.2.4 Image Stitching

Neighboring image slabs were combined using a rigid registration equiped with a simple cosine-weighted blending algorithm. Stitching two sections, A and B, together using the overlapping 0.2-cm region between the bottom of A and top of B, co-registering them together. We allowed a margin of displacement in the z-direction of 1mm (5 slices). This was done to account for potential minor inaccuracies in the mechanical device pushing the spinal cord through the MRI tube. Each displacement window ran 20 registrations between overlapping regions. The window and run with the highest mattes’ mutual information was accepted and used to stitch them together. The entirety of section B was then transformed to match section A, using the transformation matrix from the best registration between the overlapping regions. The bottom of the transformed section B was then registered to the top of the next section C in the same way as A and B were registered to each other. Repeating the process sequentially until the last section had been registered to the second last one. We initially concatenated section A and B using half from each of the overlapping regions. However, this produced strong boundary effect in the slice transitioning from section A to B. A cosine-weighted transition function between the overlapping regions was therefore employed, showing a significant reduction in boundary effects, see equations 4 and 5. Let A(x,y,z) be the intensity of section A, B(x,y,z) be the intensity of the registered section B and the overlap along the z-axis as having length N, with index k ∈ {1, 2, …, N − 1} while ω denotes the weight of the given section.

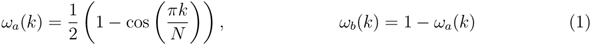

Equation 4

The blended overlap intensity is then given by I(x, y, k):

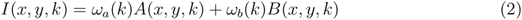

Equation 5

Where at k = 0: ω*_a_*(0) = 0, ω*_b_*(0) = 1 → *purely section B*, and at k = N−1: ω*_a_*(N−1) = 1, ω*_b_*(N − 1) = 0 → *strictly section A*. Overlapping slices in between are then a product of both weighted by the cosine transition function.

#### 5.2.5 Co-registration

We co-registered the preprocessed T2 image to an average preprocessed b0 image to minimize artificial manipulation of the dwMRI images. Negative Mattes’ mutual information was chosen as the registration cost function. Mattes’ mutual information assumes that both images have a similar sampling resolution of structures. A T2 dummy image was therefore downsampled into B0 space before being used to optimize rigid affine parameters. Co-registration was run 20 times. The run, with the lowest negative mean mutual information across the 20 runs, was then accepted. A linear registration was finally applied to the full-resolution T2 image using the chosen rigid affine parameters. All steps were executed using ANTsPY (63,67). We observed a translation of [-5.65e-3 5.79e-3 - 3.78e-2] mm in the x,y and z direction, with the co-registration requiring only minor angular corrections (< 1° across all axes).

#### 5.2.6 Bias Field Correction

Bias Field Correction: Bias field inhomogeneities are a common problem across all field strengths, especially in high-field MRI (57). They are caused by RF coil sensitivity profiles, non-uniform B_1_ transmit fields, and magnetic susceptibility effects. While they do not affect spatial geometry, bias fields distort tissue intensity profiles, creating confounding factors for intensity-based analyses such as segmentation, registration, and tissue classification. To address this, we applied an N4 bias field correction, implemented within the ANTs framework. The N4 algorithm models the bias field as a smooth, multiplicative, low-frequency field. It iteratively estimates and removes this field using a B-spline approximation to improve tissue contrast and normalize intensity across space. We first applied the N4 bias field correction to each slab individually, using the following parameters: a shrink factor of 1, with convergence set to run with [50,50,50,30] iterations at varying resolution levels. Each iteration used a tolerance of 1e-6 and a B-spline with control point spacing of 2mm in all directions, see Fig. 3C. The mean intensity profiles across all directions, i.e., z, x, and y, were flattened, indicating successful bias-field correction. This is further supported by an average 10% reduction in the coefficient of variation (CV) across the x and y directions. However, as shown in big S. 7, the mean intensity profiles of the slabs no longer appear homogeneous but have instead become dispersed due to shifts in mean intensity across the entire volume. This suggests that N4 may have mistaken true anatomical gradients, possibly from WM/GM contrast ratios or intensity drops with depth, for inhomogeneity. Leading to downstream issues with slab stitching, as co-registration expects similar intensity profiles across the overlapping section of the two slabs. We therefore choose to apply bias-field correction as the only preprocessing step after stitching, to ensure a smooth, global intensity profile across slabs. This approach also resulted in a 10% reduction in CV while minimizing boundary effects and preserving anatomical contrasts. See S. 7A & B for post-correction mean intensity profiles across the entire spinal cord and a subset of the bias fields with associated raw and corrected spinal cord images. The bias field correction procedure reduced slab boundary intensity mismatches and corrected for small intensity inhomogeneities in the axial plane.

### 5.3 Spinal Masks

#### Full Spinal Mask

All full spinal cord masks were acquired using a median filter Otsu thresholding technique (68), a basic thresholding algorithm for separating embedded distributions, i.e., different tissues or background intensities. A quick inspection of intensity histograms showed a clear multi-modal distribution across all images, thereby adhering to a core assumption of otsu thresholding. The median filter was set to use a radius of 2 voxels for dwMRI and 4 voxels for T2, with a total number of passes of 4 for both modalities. Thresholding is quite sensitive to noise, so we generated our first mask post denoising, which was henceforth used to reduce computational load and improve the performance of future preprocessing steps. A final full spinal cord mask was then generated post the remaining preprocessing steps.

#### Gray and White Matter Segmentation Mask

Training Procedure We downloaded the pre-trained model as an .onnx file, ONNX is nowadays considered the universal storage for ML models (69). Allowing cross-framework communication, i.e., train a model in one framework, e.g., PyTorch, TensorFlow, Scikit-learn, etc., and export it to ONNX format. After which, it can be loaded and run in any ML environment. We converted the .onnx file to fit our choice of modelling framework using onnx2torch (70). All training and validation of the model was done using the Pytorch framework (71). Small allterations to our data were required to make it fit with models input parameters. Namely, the original model expected an input dimensionality of (batch, 200,200, feature). Requiring cropping and downsampling of our original image resolution from 384×384 to 200×200. T2 images were interpolated using a third-order B-spline interpolation, preserving tissue boundaries while minimizing aliasing and interpolation artifacts. Associated manual or model-generated segmentation masks were resampled using nearest-neighbor interpolation to avoid class mixing. We first fine-tuned the entire parameter space, i.e., 475,329 parameters, using the pre-labeled post-mortem T2 spianl cord images from (22). This was done to bias our model towards more domain-specific information, before finally training the model on 20 slices of our own manually labeled data. We resampled all the images and labels using the procedure described above. T2 images were normalized to range between [0,1] on a slice-by-slice basis, only using foreground voxels containing spinal cord tissue for normalization. All slices either empty or containing non-target tissue were removed to avoid introducing pure noise into the training procedure, resulting in the exclusion of 140 slices. We used 80% of the data for training while keeping 10% for a validation split aimed at hyperparameter tuning, i.e., choosing an appropriate learning rate function, cost function, dropouts, and number of epochs. The final 10% test split was kept separate until the very end, providing a final unbiased evaluation of the fully trained model. We stratified the mean index of all three splits to be equal ±10%, ensuring an equal split of different spinal cord segments across the train, validation, and test splits. We chose a batch size = 15, with each batch being independently subjected to our data augmentation pipeline. At the start of each epoch, the training dataset is randomly shuffled, and the batches are drawn anew from the new shuffled order.We instantiated two networks, one for gray-matter segmentation and one for white-matter segmentation, as the pretrained network was trained on binary classification with the final layer being a sigmoid node. Both networks were trained with a maximum number of epochs = 200, but with the following stopping rule to avoid overfitting.

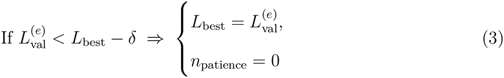

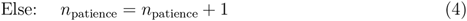

Training stops if:

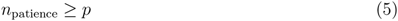

We chose δ = 10^−4^ and p = 5, i.e., training is stopped if the validation loss does not improve by at least 10^−4^ for five consecutive epochs.

#### Data Augmentation

To improve model generalizability and mitigate overfitting, we applied a combination of spatial and intensity-based data augmentations during training. Spatial augmentations were applied jointly to images and their corresponding segmentation masks to preserve anatomical correspondence. Spatial augmentation included a random horizontal flipping applied with a probability of (p = 0.5), random affine transformations with up to ±10° rotation and isotropic scaling in the range of 90%–110% (p = 0.5), elastic deformations (p = 0.2), and random cropping with padding to maintain 200×200 pixels (applied with p=0.75). These steps improve robustness and mimic possible natural anatomical variability of the spinal cord and its images, thus exposing the model to a broader range of possible spinal cords. Thereby preventing the network from overfitting to fixed specific geometric patterns leading to enhanced generalization to unseen data. Intensity-based augmentation was solely applied to the T2-weighted images, as intensity-based modulation of segmentation masks does not make sense due to their binary nature. We applied a random intensity shift to each image in the range of −0.3 to +0.3. This operation slightly brightens or darkens the image uniformly across all pixels. This is particularly useful in our case, as the Multiple Overlapping Thin Slab Acquisition procedure creates boundary effects observed as intensity shifts between slabs. But also serves as generally useful due to different MRI scanners having subtle changes in signal intensity caused by differences in hardware, coil sensitivity, or acquisition sequences. By combining geometric transformations with intensity perturbations, the augmentation pipeline ensures that the network encounters a broader distribution of plausible inputs, improving its ability to generalize to unseen subjects, field strengths, and vendors.

#### Segmentation Results

All training took place on an ASUS RTX 5060ti GPU, utilizing parallel processing with CUDA-13 drivers. Our gray-matter network reached the early stopping rule after 77epochs with a training time of 188 minutes. Reaching a training loss = 0.030 and a validation loss = 0.028, with an AUC = 0.995 and a TPR = 97.53%. Our model quickly transferred its knowledge to the new image modality after a few epochs. The full results of our gray-matter model convergence and training procedure are shown in Fig. 6 A-D. The white-matter network reached the early stopping rule after 72 epochs, with a training time of 176 minutes. Reaching a training loss = 0.015 and a validation loss = 0.014, with an AUC = 0.997 and a TPR = 98.72%. We conducted hyperparameter optimization using the validation split, including finding the optimal quantizer threshold for binarizing our models’ probabilistic predictions. The quantizer threshold was scored using the Receiver Operating Characteristic (ROC) curve, Matthews correlation coefficient (MCC), and F1 score. We manually varied the threshold between 0 and 1 by increments of 0.01, looking for the argmax value of our three metrics. A majority vote chose the final threshold between the three metrics. Yielding the respective cut-offs for white-matter and gray-matter, δ*_W M_* = 0.19 & δ*_GM_* = 0.23. With values below the cut-off being labeled as off-target and values above as either gray-matter or white-matter. A final test of the model was done on the test split withheld from training and hyperparameter optimization. The model generalized well to the unseen test-split, illustrated by achieving a DiceScore*_GM_* = 0.9751 & DiceScore*_W M_* = 0.9867, see S. 8D & H. A post-processing binary dilation of the gray-matter segmentation was applied to fill in small holes and the thin gray-commissure line. We combined the two segmentation masks by embedding the white-matter mask on top of the gray-matter mask. Causing any voxels potentially classified as both white matter and gray matter to end up as white matter. Reducing the number of potential false positive gray matter classification in white matter regions as a consequence of the binary dilation of both individual masks.

#### Spinal Segment Parcellation

We divided the longitudinal axis of the spinal cord into its spinal segments (C1, C2 … S5) based on the meta-study of (36). They reported the relative length of each spinal segment based on several studies on human spinal cord morphology. We used their population average to define the upper and lower bounds, delimitating all the spinal segment from C1-S5.

#### B-Matrix Compatibility

A common yet simple and detrimental mistake in many DW-MRI pipelines is B-matrix incompatibility (72). The B-matrix or B-tensor holds information regarding the applied diffusion encoding gradients, i.e., the diffusion gradient orientation and strength. The B-matrix is what links the image and the estimation of diffusion models and representations. Different scanner manufacturers, file formats, and software versions save and expect the B-matrix in different ways, leading to permutations and flips of directionality. Causing a potential mismatch between the coordinate systems of the B-matrix and imaging data, where lateral orientations suddenly become anterior-posterior. B-matrix incompatibilities are especially crucial if the analysis involves rotational variance analyses such as fiber tractography or fODF. But often goes unnoticed when solely analyzing rotationally invariant scalar maps such as FA, MD, etc. We therefore manually inspected the diffusion tensors’ (DT) first eigenvectors to reveal axes permutations and gradient flips (72). We discovered a permutation of the x and y axes, which would have otherwise resulted in commissural crossing-fibers appearing to be dorsal-ventral orientated. A simple fix was implemented by transposing the x and y axes, leading to compatibility between our B-matrix and image dimensions.

### 5.4 dwMRI Analysis

#### 5.4.1 Model 1: DTI

Diffusion tensor imaging (DTI) is one of the most established techniques for characterizing diffusion-weighted MRI (dwMRI) data. It models water diffusion within each voxel as a three-dimensional Gaussian process, thereby representing the diffusion profile as a second-order tensor (D). Fiber orientations can be assessed by modelling diffusion as a Gaussian distribution, estimating quantitative parameters such as mean diffusivity (MD), fractional anisotropy (FA), and apparent diffusivity (AD). Providing a popular means to investigate tissue microstructure (73–75). However, the parameter estimates of the DTI model do not relate to specific features within tissue microstructure and are consequently sensitive to multiple structural compartments concurrently (33). We nonetheless included it in our analysis due to being the most widely used model for dwMRI data. The diffusion matrix of the DTI model *D*is defined as a 3×3 symmetric positive-definite matrix and calculated by solving a linear least-squares fit of the Stejskal–Tanner equation for every voxel (76, 77).

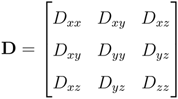

Eigen-decomposition of the **D** matrix produces eigenvalues λ_1_, λ_2_, λ_3_ sorted by principal diffusivity, with associated eigenvectors e⃗_1_, e⃗_2_, e⃗_3_, giving us the principal direction of diffusion. From here, we can compute the aforementioned biologically interpretable scalar metrics, i.e., FA, MD, AD, and RD for biological interpretation. Mean diffusivity (MD) reflects the average magnitude of water diffusion, regardless of direction:

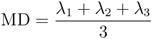

Axial diffusivity (AD) quantifies the diffusion along the principal axis:

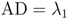

Radial diffusivity (RD) measures the average diffusion perpendicular to the main fiber. axis

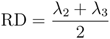

FA quantifies the degree of directional dependence of water diffusion within a voxel and is bounded between [0, 1], with 0 indicating isotropic diffusion (equal diffusion in all directions) and 1 indicating complete anisotropic diffusion.

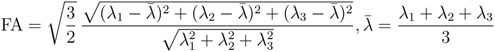

DTI models the MR signal as a single, Gaussian diffusion process per voxel, with S(b, **g**) = S_0_ exp −b **g**^⊤^**D g**, where b = γ^2^G^2^δ^2^ Δ − *δ*. However, approximating the diffusion directions using a second-order tensor brings several fundamental limitations when our goal is to map fine microstructure. tissue is comprised of multiple compartments with distinct micro-environments of water behavior due to membranes, morphology, and macromolecular crowding. Each compartment with their own unique restriction, hindrance, and exchange parameters. So, when tissue contains multiple compartments such as membranes, organelles, axonal walls, or microstructural disorder that constrain motion, it results in molecules bouncing off boundaries, getting funnelled along axons, or switching between compartments (78).

The resulting displacement distribution is therefore not Gaussian as assumed by the DTI model, and as seen for unrestricted Brownian motion (79). This is especially a problem at higher b values with longer diffusion times, like the ones we are employing. Providing more diffusion-weighting, which is particularly useful for sensing slow-moving water molecules and smaller diffusion distances, but also further probes the signal from water molecules that are more restricted in their movement due to cellular compartments and barriers. Consequently, it produces diffusion with a non-Gaussian displacement distribution. Another problem with DTI is intra-voxel heterogeneity. Voxels contain multiple fibers with varying orientations, giving rise to branching, crossing, or fanning fibers. The DTI effectively only recognizes the second moment of orientation and collapses everything else into one tensor, even when a voxel contains multiple fiber populations or orientation dispersion. This means that the tensor model represents one independent, dominant direction per voxel, resulting in estimated orientations that are ambiguous or misleading in voxels with more complex fiber structures (80, 81). For example, branching, fanning, and crossing fibers with different orientation dispersions can all produce the same tensor diffusion, despite having different morphologies. Unidirectional white matter tracts are thus solvable by DTI, but the intricate structure of gray matter remains too complex for it to resolve. It implies that DTI is excellent for macroscopic anisotropy and unidirectional tract orientation, but is not designed to resolve fine-scale microstructure. In our case, it reveals macroscopic anisotropic differences across the spinal segments, specifically that the diffusion profile of the thoracic and sacral segments diverges significantly from that of the rest of the spinal segments.

#### 5.4.2 Model 2: Neurite Orientation Dispersion and Density Imaging

Neurite Orientation Dispersion and Density Imaging (NODDI) is a multi-compartment diffusion MRI model designed to probe the microstructural organization of neural tissue, far exceeding the possibilities of conventional diffusion tensor imaging (37). As mentioned, DTI is limited by its base assumption of Gaussian diffusion and a single fiber orientation per voxel. Whereas NODDI decomposes the diffusion signal into three biophysically meaningful components: (1) an intracellular neurite compartment, representing restricted diffusion within axons and dendrites, (2) an extracellular compartment, capturing hindered diffusion in the space surrounding neurites, and (3) an isotropic compartment, modelling free water such as cerebrospinal fluid or fixative buffer. The dMRI signal 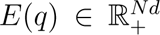, where q denotes the diffusion gradient, is described by the hierarchical model in equation 6, where 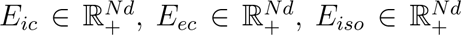 are the normalized dMRI signals of the intra-cellular, extra-cellular, and isotropic compartments. With v*_ic_*, v*_ec_* & v*_iso_* corresponding to the volume fractions, constricted by v*_ec_* = 1 − v*_ic_*, E*_ic_* and E*_ec_* adopt the orientation-dispersed cylinder model introduced in (zhang, 2011), which are based on a Watson probability distribution W (µ, κ), for modelling orientations on the unit sphere S^2^. With all orientations being rotationally symmetric about µ ∈ S^2^ and whose concentration parameter κ controls the amount of dispersion around it.

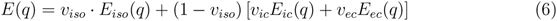

From this model, we can derive more detailed voxel-wise maps of microstructure through its fitted parameters. Namely, neurite density index (NDI) bound between 0 and 1, with one representing tightly packed axons and dendrites and zero being indicative of a high v*_iso_*. Orientation dispersion index (ODI) is bound between 0 and 1 and calculated as 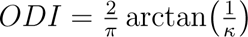. One indicates a high dispersion rate in all directions, whereas low values signal a concentrated and coherent diffusion direction. Lastly, we have the free water fraction (FWF) metric accounting for confounding diffusion arising due to the partial volume effect. These metrics provide complementary information about the neural microstructure, with NDI reflecting the density of axonal and dendritic processes, ODI describing the angular coherence of neurite orientations, and FWF quantifying the contribution of isotropic free water. Intuitively, a change in FA derived from a DTI model can be a result of two main microstructural changes. (1) A decrease in the dispersion of neurite projections, corresponding to a homogenous principled direction. Water would thus only diffuse along the gradient of the restricted neurite projection, resulting in the following model parameters: a lower ODI due to decreased dispersion, and a stable NDI, as no changes in axonal and dendritic packing had been observed, which in combination yields a higher FA. (2) The other way we could expect the FA to increase is due to an increase in axonal and dendritic packing density. Closer-packed neurites would result in limited diffusion, due to increased mass of boundaries and hindrance of diffusion. This would be reflected by an increase in FA despite no changes to the orientation distribution. We utilized the Accelerated Microstructure Imaging via Convex Optimization (AMICO) (82), which reformulates the NODDI model fitting as a convex optimization problem, thereby reducing the fitting time from days to hours, while still maintaining the accuracy and reproducibility of the original fitting procedure.

#### 5.4.3 Model 3: CSD

Principled direction does not represent neurite orientations equally for all voxels. Voxels with a low ODI are sufficiently described by the principled direction, similar to how the mean is a good model of a distribution if the standard deviation is low. Highly parallel white matter tracts can therefore sufficiently be depicted with this direction-encoded colored FA maps, but the more nuanced gray matter needs something more complex representation to accurately depict existing micro structural variations. We therefore acquire a descriptive measure that captures the heterogeneity of directionality when variability is high. We achieve this with a spherical deconvolution model, that deals with nonsingular fiber orientation distribution, such as crossing-fibers. Spherical deconvolution is a particularly appealing method for our HARDI data as it can provide estimates of the whole fiber orientation distribution function (fODF) across entire hemisphere per voxel (83). Making it ideal for our need to estimate intra-voxel fiber orientations in gray matter structures. Practiacally spherical deconvolution models work by fitting the diffusion-weighted signal on the unit sphere as a spherical convolution of a single highly coherent fiber response function with an unknown fiber-orientation distribution (FOD). Relying on the principle that the diffusion weighted signal originating from the different fiber populations in a voxel is given by the spherical convolution of the response function with the fODF. The desired fODF can thus be estimated by performing the spherical deconvolution of the response function from the measured diffusion weighted signal. We used a linear basis of spherical harmonics (SH) to achieve a reduced compact representation of the spherical function (83). First step is to estimate the coefficients of the single-fiber response function from the diffusion weighted signal. This is done by identifying voxels deemed to contain a single coherent fiber orientation, corresponding to a single compartment response function, and fitting a basis function to it. A spherical convolution matrix can then be formed, relating the vector of unknown fODF coefficients to the vector of diffusion weighted signal intensities, and the spherical deconvolution operation can be cast as a linear least-squares problem. However, the estimation of the singular response function still only comprises a single-compartment model, which does not overcome the partial-volume effect. So, we implement a multi-tissue extension of the spherical deconvolution model, further decomposing the signal into a white-matter FOD and isotropic compartments characteristic of gray matter and cerebrospinal fluid, similar to the NODDI model. By fitting a response function individually for each compartment, in our case, white matter, gray matter, and the buffer used for fixation. Thus, mitigating the partial-volume effects leading to an accurate fODF reflecting microstructural changes in the gray matter hinted at by the DTI analysis. For this we based our model on the RUMBA-SD algorithm equiped with spatial regularization (38) to generate a convolution kernel mapping of the fODF on a symmetric half-sphere with 724 equally spaced points of the recorded data. The fODFs are then estimated using an iterative, maximum likelihood estimation algorithm adapted from Richardson-Lucy (RL) deconvolution (84)). Fitting took 925minutes on an Intel(R) Core(TM) Ultra 7 255HX. We represent the fODF by 3D glyphs in the FURY framework in python.

## 6 Microscopy

In this section we outline a detailed version of the microscopy pipeline, including everything from the data-collection, preprocessing and the various analytical approaches utilized throughout the study.

### 6.1 Data Acquisition

The dura mater was removed, and the spinal cord was segmented transversally into cervical, thoracic, lumbar, and sacral regions. Each region was further sectioned into 50 µm transverse slices using a microtome, and sections were stored in 1X PBS containing 0.2% sodium azide. For immunohistochemistry free-floating 50 µm spinal cord cross-sections from all spinal regions were washed three times in 1X PBS every 5 min. Sections were then blocked in blocking solution for 2 h at room temperature as previously described (85), followed by three washes in 1X PBS. Subsequently, sections were incubated with primary antibodies diluted in antibody solution overnight at 4°C. The following primary antibodies were used: anti-SMI32 (mouse monoclonal, 1:1000 dilution; Chemicon, Millipore, Milan, Italy); anti-NeuN (rabbit monoclonal, 1:500, abcam AB177487; anti-GFAP (goat polyclonal, 1:500, Invitrogen, PA5-143587), anti-S100 (rabbit polyclonal, 1:1000, DAKO, Glostrup Denmark, Z0311); and anti-ChAT (goat polyclonal, 1:400, EMD Millipore AB144P). The following day, sections were washed three times in 1X PBS (10 min each) with gentle shaking. Secondary antibodies (Alexa-Fluor 647 donkey anti-mouse, 1:1000 dilution (KU, A-31571); Alexa-Fluor 488 donkey anti-rabbit (Invitrogen, A32790), 1:500; Alexa-Fluor 594 donkey anti-goat, 1:500 and 1:1000 (abcam, ab150132) were added to 500 µl of antibody solution together with Hoechst for nuclei staining, and the tissue slices were kept in the dark for 2 h at room temperature. Later, the sections were washed in 1X PBS 3 times in the dark, followed by a fast wash with distilled water at room temperature. Sections were air-dried and mounted on the glass slides for imaging. Zeiss Cell Observer spinning disc microscope was used for imaging with a widefield module. A 20X objective with a 0.8 numerical aperture was used with a working distance of 2.55 mm. Light sources with a diode laser of 405 nm, an argon laser of 488 nm and 561 nm and a HeNe laser of 635 nm were used. Emission filter cubes with wavelengths of BP450/50 for Hoechst, BP525/50 for AlexaFluor 488, BP562/45 for Alexafluor 555, BP629/62 for AlexaFluor 647 were used. During imaging, approximately 60 - 80 z-stacks were imaged per segment. Large fractions of the 3D volumes were out of focus, most likely as a result of uneven slicing, resulting in low contrast and SNR over extended regions, leading to hampered robust phase-correlation estimation for stitching. Resulting in large parts of the 3D volume being unusable, roughly 8 to 10 z-slices per z-stack had more than 80% of the axial plane in focus. This made robust estimations of longitudinal projection impossible and we thus had to limit our analysis to the 2D plane.

### 6.2 Preprocessing

#### 6.2.1 Flat Field Correction

Our fluorescence images suffered from a heavy degree of vignetting, these shading artifacts manifest as smooth, low-frequency intensity gradients that, typically observed as an decreasing illumination gradient as we move distal from the optical center. If left uncorrected would it introduce systematic bias into downstream quantitative analyses and produces strong boundary artifact among adjacent slabs post image stitching. To correct for these artifacts, we employed BaSiC (Background and Shading Correction), a retrospective correction method developed by (86), based on low-rank and sparse decomposition of an image. The method estimates a smooth multiplicative shading field and an additive background component directly from the acquired images, thereby requiring no reference flat-field images or manual parameter tuning to correct the illumination artifacts. All images were processed individually before image stitching was achieved.

### 6.3 Stitching

Acquired image tiles were stitched together using BigStitcher (87) within Fiji. The raw tiles were imported as a BigStitcher dataset and resaved to the multi-resolution BigDataViewer HDF5 format, with initial tile positions taken from the meta data on recorded stage coordinates. Pairwise translations between overlapping tiles were estimated by phase correlation, computing shifts on the overlap regions with default downsampling. The resulting pairwise links were filtered by cross-correlation coefficient, retaining only links with r ≥ 0.7 and discarding shifts exceeding the expected overlap, so that unreliable pairs were excluded from the alignment. Aligned tiles were fused into a single volume over the bounding box of the registered dataset using weighted linear blending to suppress seam artifacts in the overlap regions

### 6.4 Denoising

Fluorescence microscopy images are inherently subject to noise, i.e., photon shot noise, detector read noise, and background fluorescence just to name a few. A low SNR obscures fine structural details and hampers our downstream quantitative analysis of the fiber orientations. Applying N2V prior to structure tensor analysis ensured that observed spatial variations in fiber orientation reflected true underlying microstructural organization rather than noise-driven gradients, improving the reliability of orientation estimates particularly for regions with low signal intensity. To address this, we employed Noise2Void (N2V), a self-supervised deep-learning denoising framework by (39), producing state of the art denoising directly from single noisy images without the need of a clean version. We trained a version of N2V for each of the images using a patch size = 512×512, a ROI size = 8. Training lasted for 100 epochs with 50 number of steps per epoch and a batch size of 15. The model weights for the epoch with the lowest score within the last 10 epochs was used to denoise the full image.

### 6.5 Microscopy Analysis

Local fibre orientation and structural anisotropy were estimated from fluorescence images using a 2D structure tensor framework. For each spinal region, the best focal plane was selected prior to analysis. The structure tensor **S** was computed at each pixel using a gradient smoothing scale σ= 2.5 px and an integration scale ρ = 10 px; σ controls the scale at which image gradients are estimated, while ρ defines the ‘neighbourhood over which orientation information is pooled, suppressing noise whilst preserving mesoscale fibre structure. The structure tensor analysis returns a matrix of 3-by-3 matrix capturing the local orientation around a point in space. For each point in the volume, *V*, we can compute the structure tensor as;

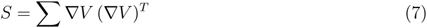

*V* is the gradient of and the integration is bounded by the neighborhood around the point set by ρ. We utilize a Gaussian kernel window for integration and a Gaussian derivative kernel for computing the gradients. Which gives us the following;

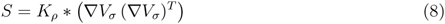

σ the parameter is the standard deviation of the Gaussian derivative kernel used for computing the gradient ∇V*_σ_*, and the parameter ρ is the standard derivation of the Gaussian kernel K*_ρ_* used for integration. Fiber-like structures and their orientation corresponds to the direction of least change in intensity. This can be formulated as minimizing the quantity **u***^T^* S**u**, where **u** is a unit vector. The vector that minimizes this expression indicates the direction of least change. We obtain this direction analytically through the eigen-decomposition of S. Since S is symmetric and positive semi-definite, it has two non-negative eigenvalues with corresponding orthogonal eigenvectors. The principal orientation, i.e., the direction of least change, is given by the eigenvector **v** associated with the smallest eigenvalue. In fiber-like structures, this smallest eigenvalue is typically much smaller than the other two, reflecting strong directional consistency. In our analysis, however, we do not explicitly use the eigenvalues. Instead, we focus on the eigenvector, which encodes orientation but not direction and MRI comparable metrics such as anisotropy etc. To quantify local anisotropy of the orientation distribution two measures were computed:

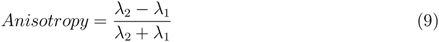

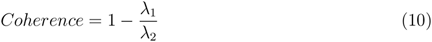

where λ_1_ ≤ λ_2_. Pixels with undefined values (e.g., due to division by zero) were excluded from further analysis. The local orientation was derived from the eigenvector corresponding to the smallest eigenvalue. Specifically, we calculated the orientation angle θ as;

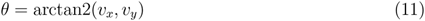

where (v*_x_*, v*_y_*) are the components of the eigenvector. To ensure consistency across slices, a global rotation offset (derived from grey matter alignment) was applied to the orientation angles. The resulting angles were wrapped to the interval [0, π] to account for orientation ambiguity. We converted the pixel-wise distributions of θ to a classical RGBA colour mapping. Fibres labelled red correspond to θ = 0^◦^, i.e., lateral orientations, whereas light blue is associated with a strictly dorsal–ventral orientation, i.e., θ = 90^◦^. We weighted the base colour mapping with anisotropy by setting the alpha channel equal to the anisotropy value. High-anisotropy fibre bundles are thus rendered with more vivid colours, whereas low-anisotropy regions produce a dimmed appearance.

## 7 Statistical Tests

### 7.1 dwMRI

#### 7.1.1 FA & ODI Cohen’s *d*

Quantification of our two diffusion metrics, FA and ODI, across spinal cord regions and tissue types was estimated by pairwise effect sizes using Cohen’s *d* for all combinations of region–tissue groups, i.e., cervical GM/WM, thoracic GM/WM, lumbar GM/WM, and sacral GM/WM. Classical null-hypothesis significance tests (e.g. Shapiro–Wilk, *t*-tests) were considered uninformative, as trivially small differences would reach statistical significance given the large sample sizes of n > 5 million per group. Cohen’s *d* was therefore preferred as a direct measure of practical effect magnitude. For each pairwise comparison, Cohen’s *d* was computed as the mean difference divided by the pooled standard deviation, with 95% confidence intervals derived from the large-sample approximation of the standard error of *d*,

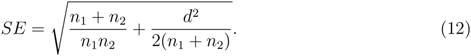

Effect sizes were interpreted according to conventional thresholds: small |d| < 0.5, medium 0.5 ≤ |d| < 0.8, and large |d| ≥ 0.8. Results were visualized as symmetric heatmaps using a diverging color scale centered at zero, with each cell annotated with the Cohen’s *d* value and its 95% confidence interval. Shown in Fig. 2.

**Spatial Autocorrelation**: Cohen’s *d* do not account for spatial autocorrelation. So we performed additional statistical tests to verify the results and their significance when controlled for spatial autocorrelation and multiple comparison. Prior to statistical modelling, spatial autocorrelation within each section was assessed using Moran’s I. For each tissue type and spinal segment combination, a distance-band spatial weights matrix was constructed using pairwise Euclidean distances. The weights matrix was row-standardised prior to computation. Statistical significance was evaluated using 499 permutations, with sections considered spatially autocorrelated at α = 0.05. To account for the hierarchical structure of the data and residual spatial dependency, a Linear Mixed Model (LMM) was fitted with FA as the outcome variable. FA values were z-score standardised prior to modelling. Fixed effects included tissue type (WM vs. GM), spinal segment, their two-way interaction, and a spatial covariance trend term. The spatial trend was derived per section by fitting a quadratic polynomial surface in the xx yy xy plane (basis: 1, x, y, x^2^, y^2^, xy) and extracting the fitted values, thereby absorbing within-section spatial structure as a fixed covariate. Spinal section (z-index) was included as a grouping factor for the random intercept. Models were fitted using Restricted Maximum Likelihood (REML) via the L-BFGS optimiser. Overall model significance was assessed by a likelihood ratio test against an intercept-only null model, refitted using maximum likelihood. Post-hoc comparisons were conducted using Tukey’s Honestly Significant Difference (HSD) test, applied across all tissue type × segment group combinations to control the familywise error rate. Comparisons were considered statistically significant at α = 0.05.

### 7.2 Microscopy

#### 7.2.1 Anisotropy

To enable spatial statistical analysis of fluorescence microscopy data acquired at 0.325 µm/pixel resolution, pixel-level anisotropy measurements were aggregated into non-overlapping square patches of 250 × 250 pixels (81.25 × 81.25 µm). For each patch, the mean anisotropy, coherency, and orientation angle were computed across all contributing pixels. Patches in which fewer than 66% of pixels belonged to the tissue mask were excluded to avoid edge artefacts and incompletely sampled regions. This aggregation reduced the dataset from approximately 1.7710^9^ pixels to a tractable number of spatially discrete observations (patches), which served as the unit of analysis in all subsequent steps. To assess whether patch-level anisotropy values exhibited spatial clustering within each tissue compartment and spinal segment, Moran’s I statistic was computed for each Segment × Tissue combination. A distance-band spatial weights matrix was constructed per group, with the neighbourhood threshold set to the 25th percentile of all pairwise inter-patch distances, ensuring each patch was connected to its nearest neighbours while remaining adaptive to local patch density. Weights were row-standardised prior to computation. Statistical significance was assessed using a permutation procedure with 499 permutations, and results were considered significant at α = 0.05. To test for differences in anisotropy across spinal segments and tissue types, an ordinary least squares (OLS) regression model was fitted with patch-level anisotropy as the outcome variable. As only a single histological section was available per spinal segment, a random intercept structure was not estimable and a fixed-effects-only model was used. Anisotropy values were z-score standardised prior to modelling. Fixed effects included tissue type (WM vs. GM), spinal segment (cervical, thoracic, lumbar, sacral), their two-way interaction, and a spatial trend covariate. The spatial trend was derived per Segment × Tissue section by fitting a second-order polynomial surface in the spatial coordinates, with basis terms {1, x, y, x², y², xy}, using ordinary least squares. The fitted values of this surface were included as a fixed covariate to absorb within-section spatial structure. Given evidence of spatial autocorrelation in patch-level residuals (Moran’s I), heteroscedasticity and autocorrelation consistent (HAC) standard errors were computed using a Newey-West sandwich estimator with band-width set to √n where n is the number of patches per comparison. All inference on model coefficients was based on HAC-corrected standard errors. Model fit was evaluated against an intercept-only null model using a likelihood ratio test, with the test statistic compared to a χ^2^ distribution. **Post-hoc Comparisons** Two complementary post-hoc strategies were employed. First, Tukey’s Honestly Significant Difference (HSD) test was applied across all pairwise combinations of Segment × Tissue groups, controlling the familywise error rate at α = 0.05. Each pairwise comparison was implemented as a two-sample OLS regression with Newey-West HAC standard errors, with bandwidth set to √n, ensuring consistency with the main model. The resulting p-values were corrected for multiple comparisons using the Ben-jamini–Hochberg false discovery rate (FDR) procedure, with significance threshold α = 0.05. All analyses were performed in Python using the statsmodels, libpysal, and esda libraries.

## 8 Supplementary Figures

**Figure 7:**
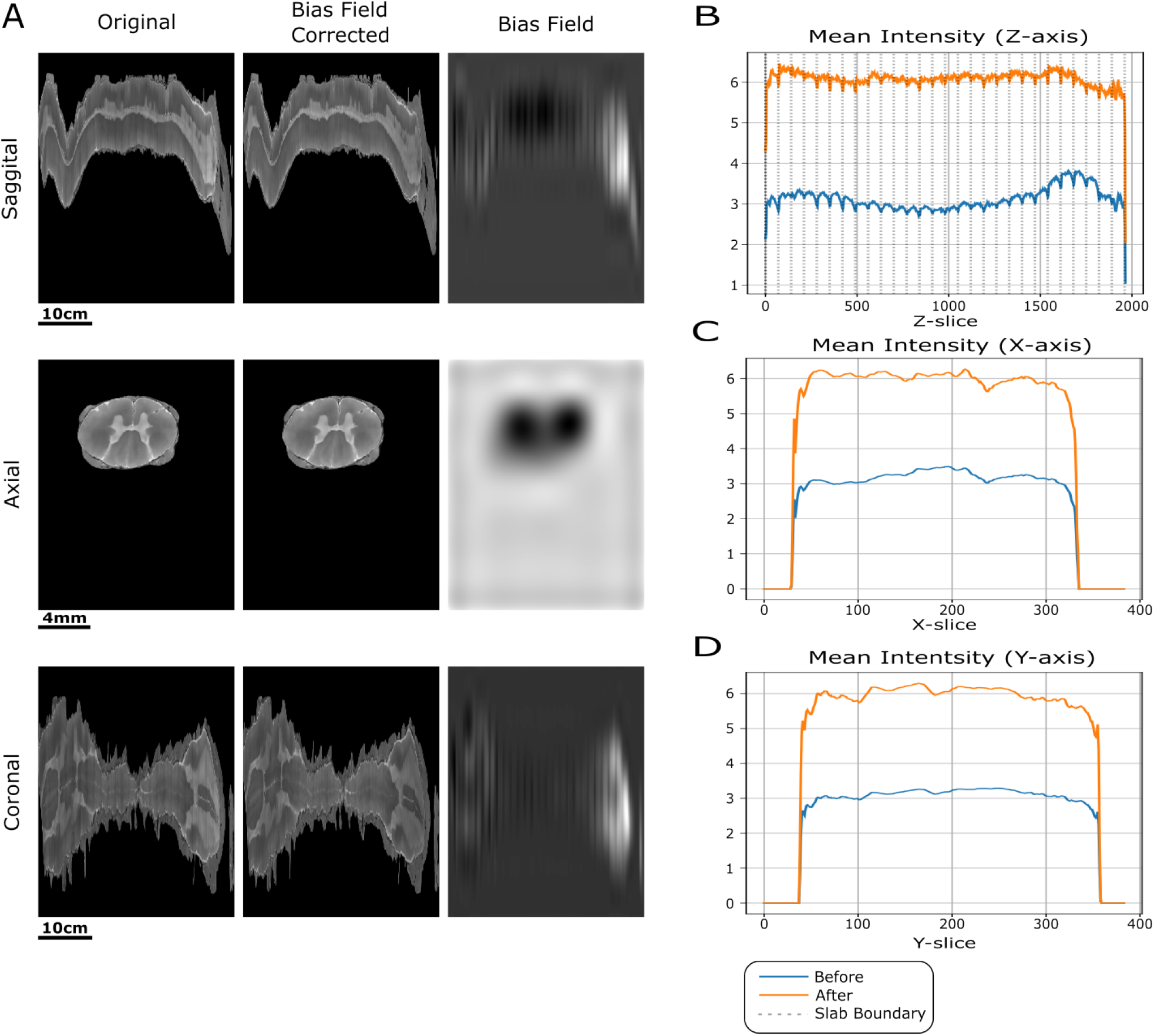
A) Depicts a saggital, axial and coronal view of the, from left to right; original image, the bias field corrected image and the bias field itself. B, C & D) contains the intensity profiles before and after bias field correction.

**Figure 8:**
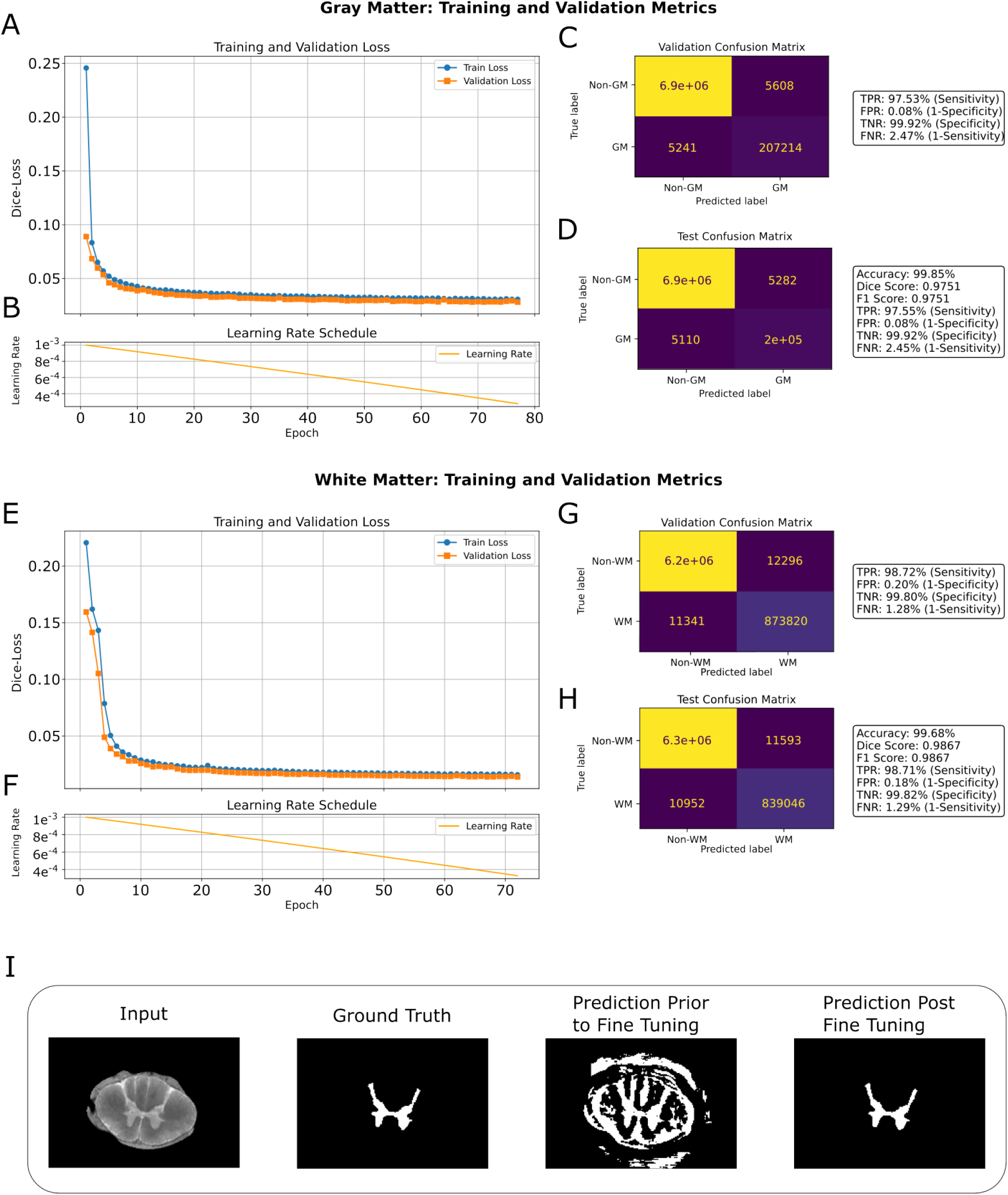
Training, validation, and test performance of the deep learning segmentation model for gray matter (GM) and white matter (WM). Gray matter metrics: (A) Training and validation Dice-loss curves over 80 epochs. (B) Learning rate scheduler. (C) Validation confusion matrix, (D) Test confusion matrix. White Matter metrics: (E) Training and validation Dice-loss curves over ~75 epochs, (F) Learning rate scheduler (G) Validation confusion matrix, (H) Test confusion matrix. (I) Segmentation examples of GM network: ground truth mask, model prediction prior to fine-tuning, and model prediction post fine-tuning.

**Figure 9:**
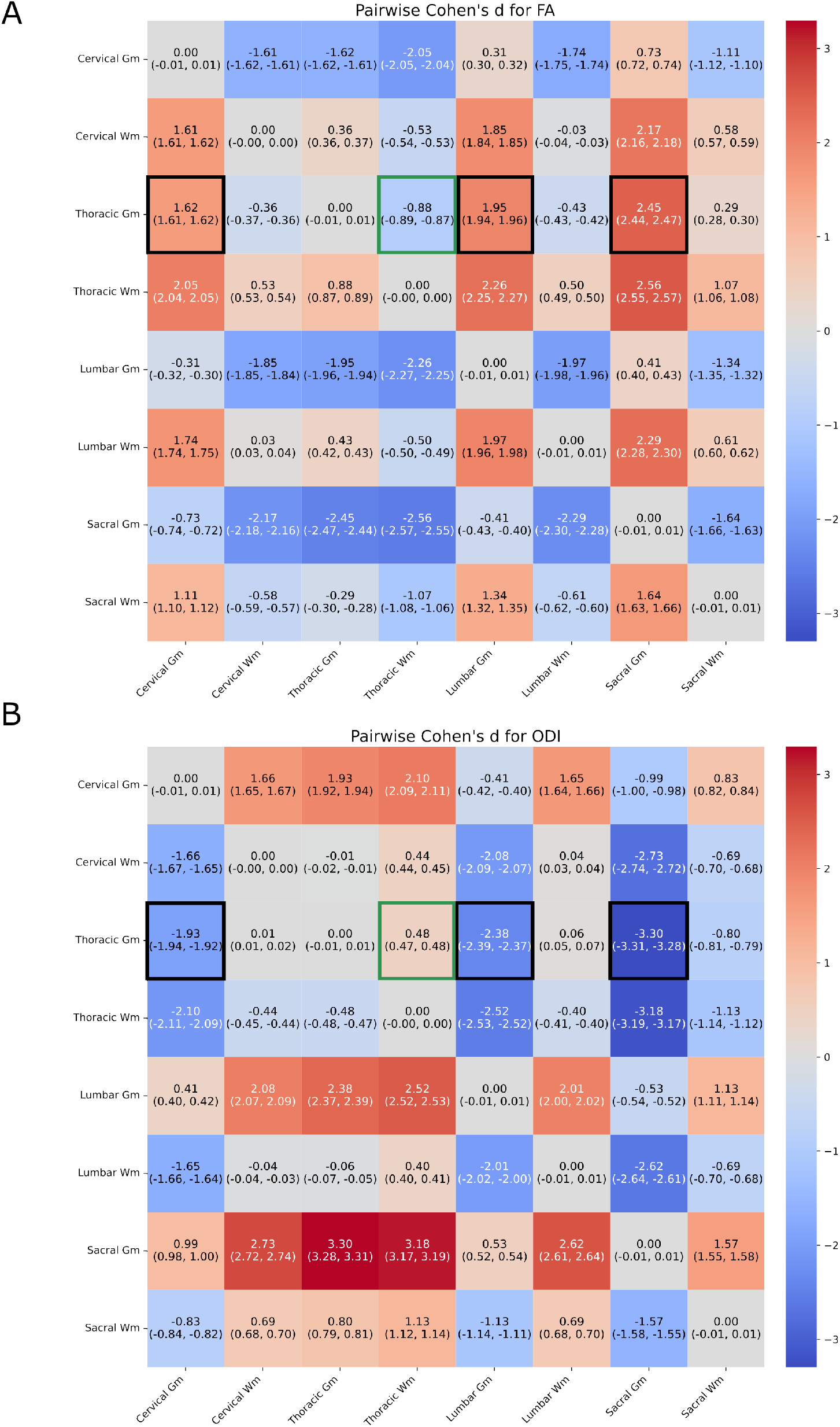
Full scale heatmap of Cohen’s D of FA in (A) and ODI in (B) with all combinatorics of tissue type and segment.

**Figure 10:**
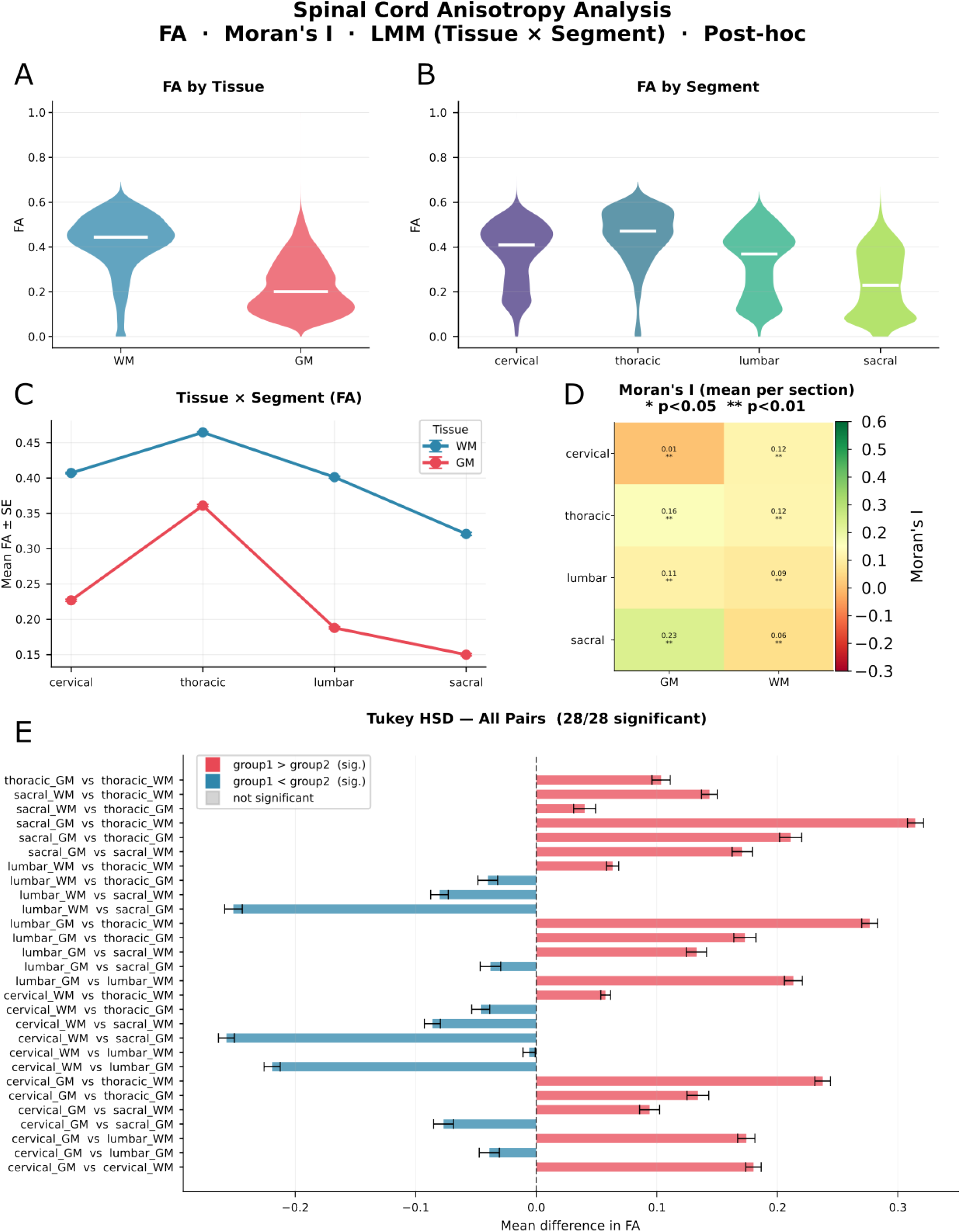
Overview of in depth statisical tests performed on dwMRI derived fractional ansiotropy (FA). A & B) Shows the mean FA across segment and tissue type. C) depicts FA stratified by tissue type, segment and their interaction. D) Moran’s I a measure of spatial autocorrelation. E) Depicts all post-hoc comparisons from the linear mixed effect model FAT̃issue x Segment, with standard errors corrected for multiple comparison.

**Figure 11:**
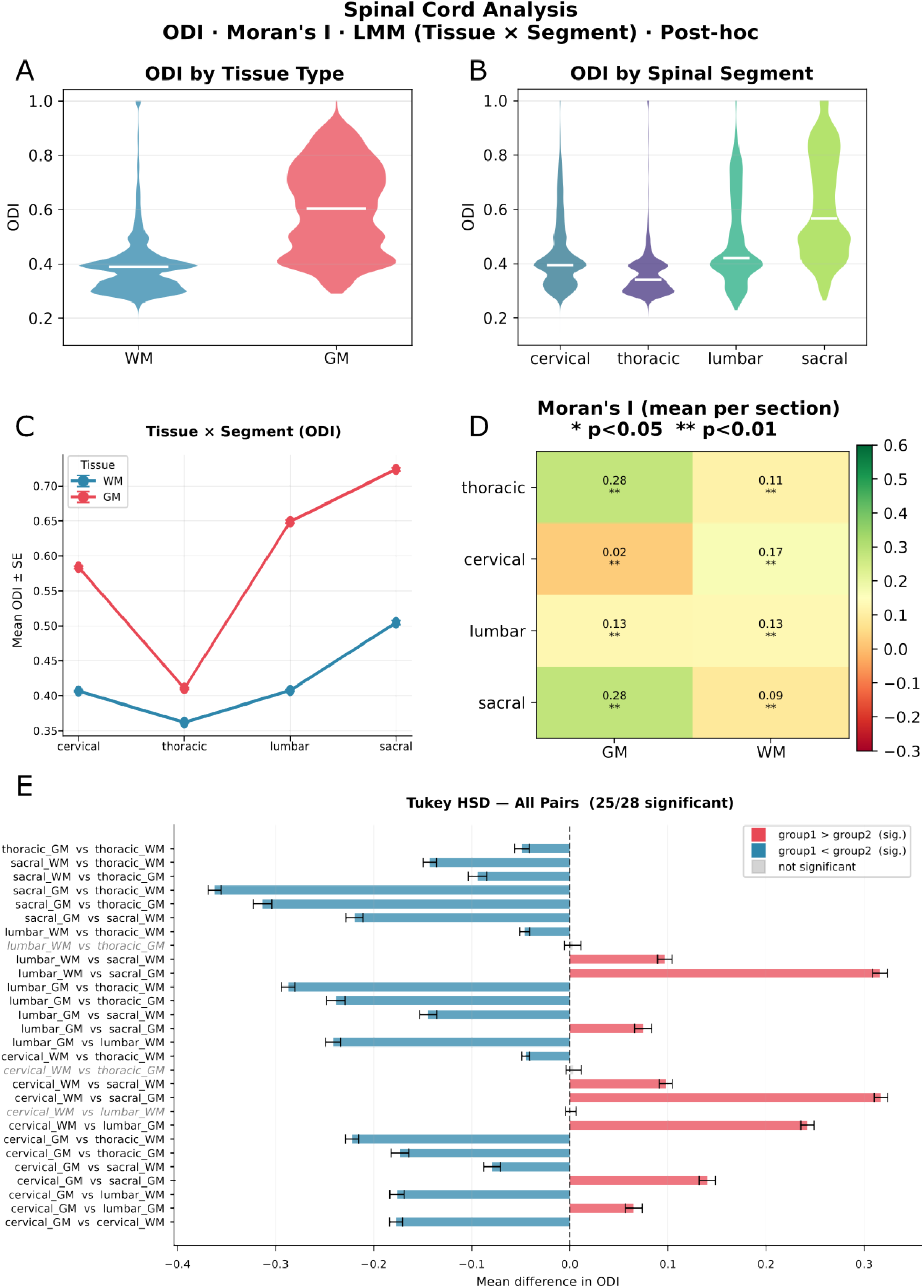
Overview of in depth statisical tests performed on dwMRI derived ODI. A & B) Shows the mean ODI across segment and tissue type. C) depicts ODI stratified by tissue type, segment and their interaction. D) Moran’s I a measure of spatial autocorrelation. E) Depicts all post-hoc comparisons from the linear mixed effect model ODIT̃issue x Segment, with standard errors corrected for multiple comparison.

**Figure 12:**
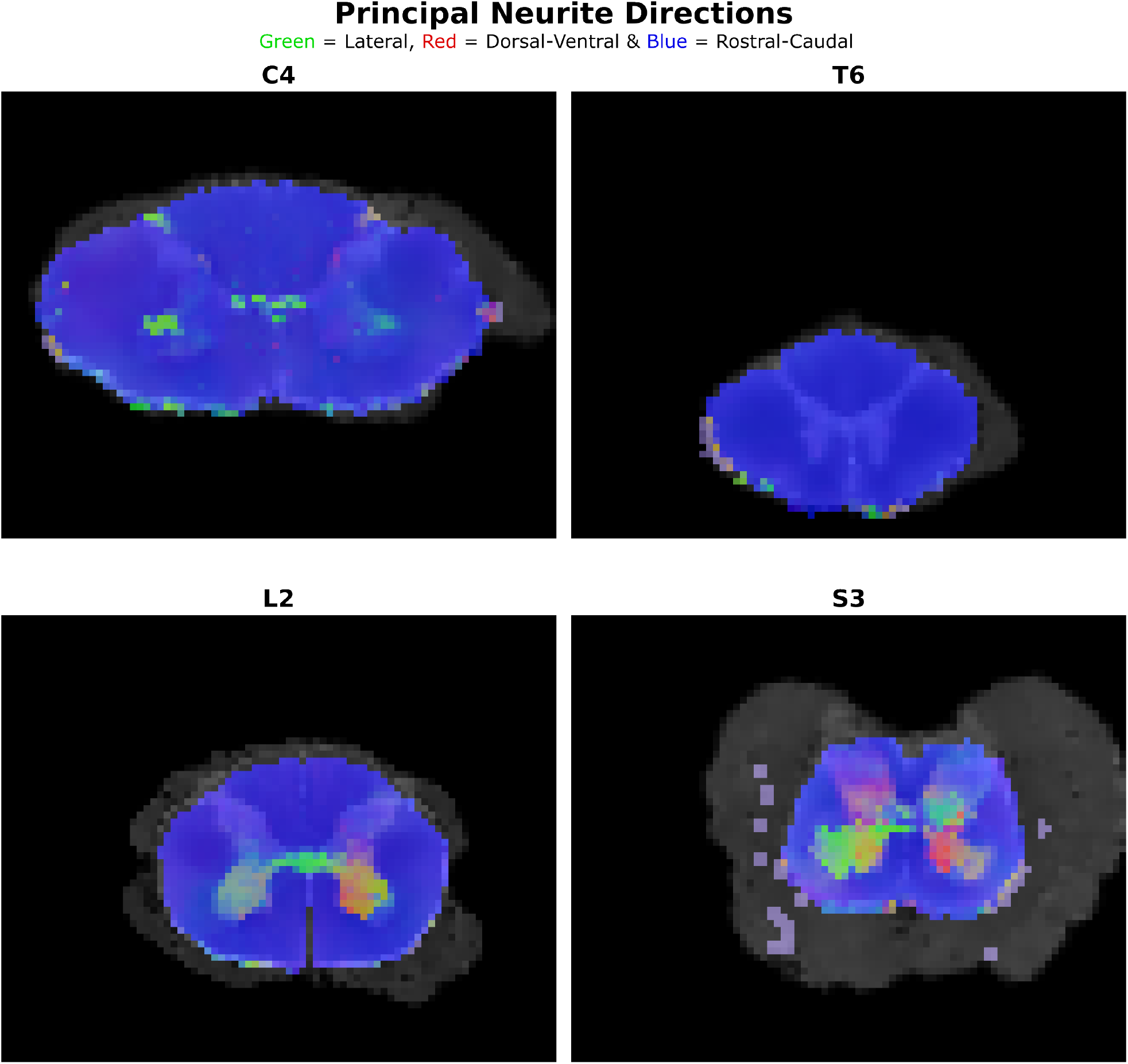
Principal neurite orientations dervied from NODDI model. Blue indicates a pre-dominant rostro-caudal direction, green a lateral and red a dorsal ventral.

**Figure 13:**
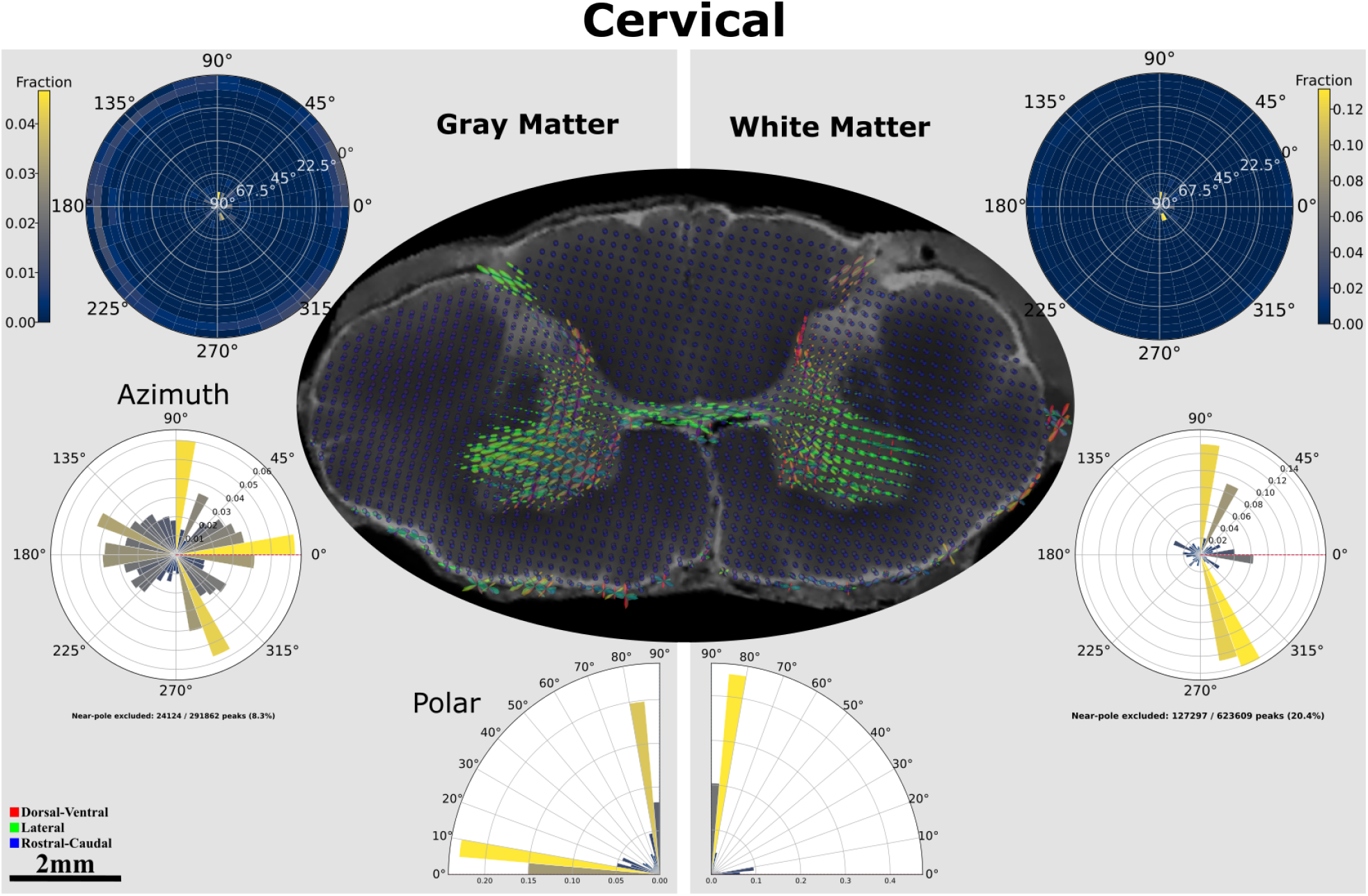
Quantifiation of cervical fODF profiles on a hemisphere stratified by tissue type. Upper left and right depicts a density plot of a flattend hemisphere. With the radius i.e, the angles depicted in white denoting polar angles, and azimuth angles by the circular angles. Forming a fully 3d representation of the fODF rather than the individual polar and azimuth distributions shown below.

**Figure 14:**
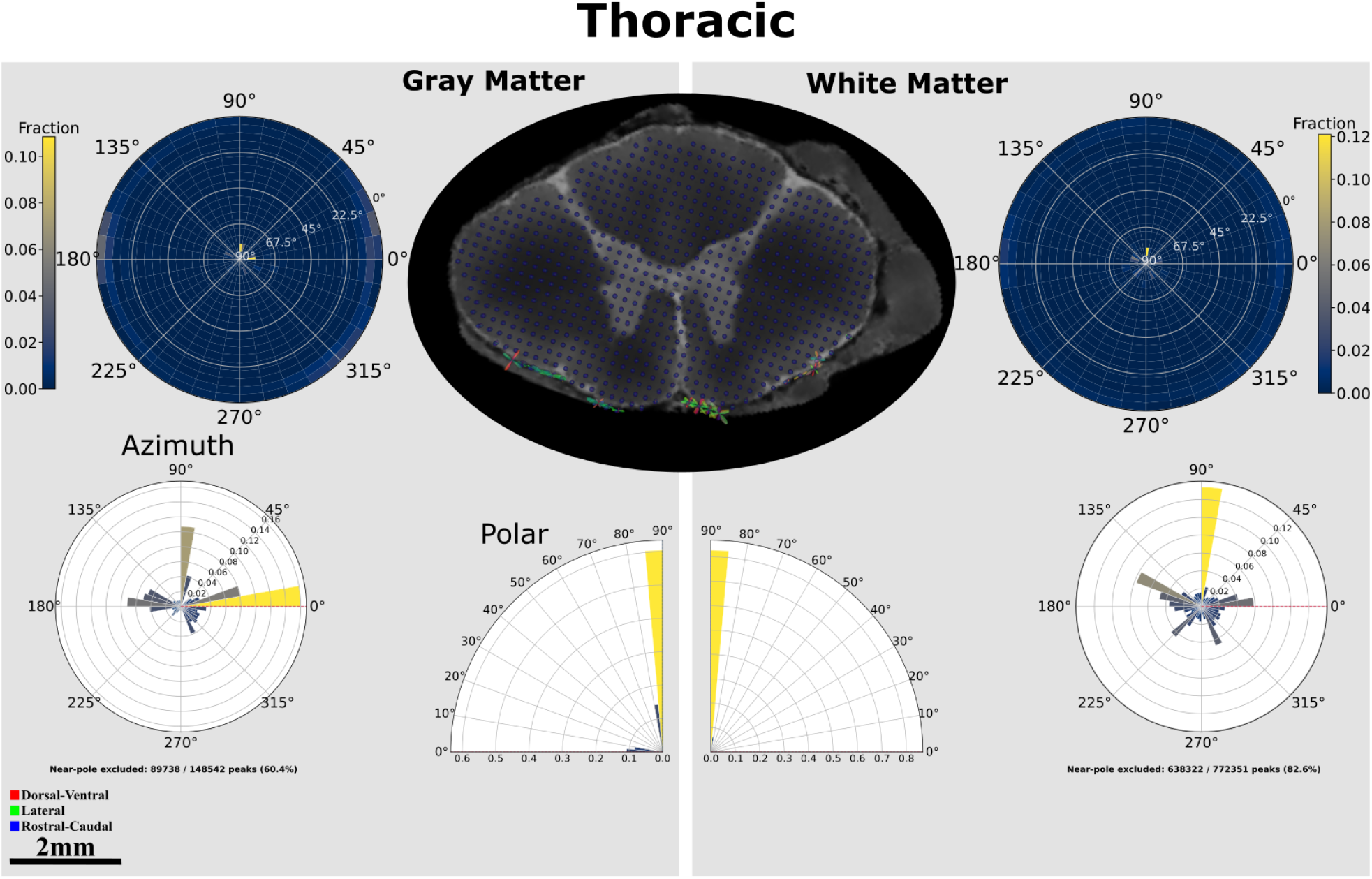
Quantifiation of thoracic fODF profiles on a hemisphere stratified by tissue type. Upper left and right depicts a density plot of a flattend hemisphere. With the radius i.e, the angles depicted in white denoting polar angles, and azimuth angles by the circular angles. Forming a fully 3d representation of the fODF rather than the individual polar and azimuth distributions shown below.

**Figure 15:**
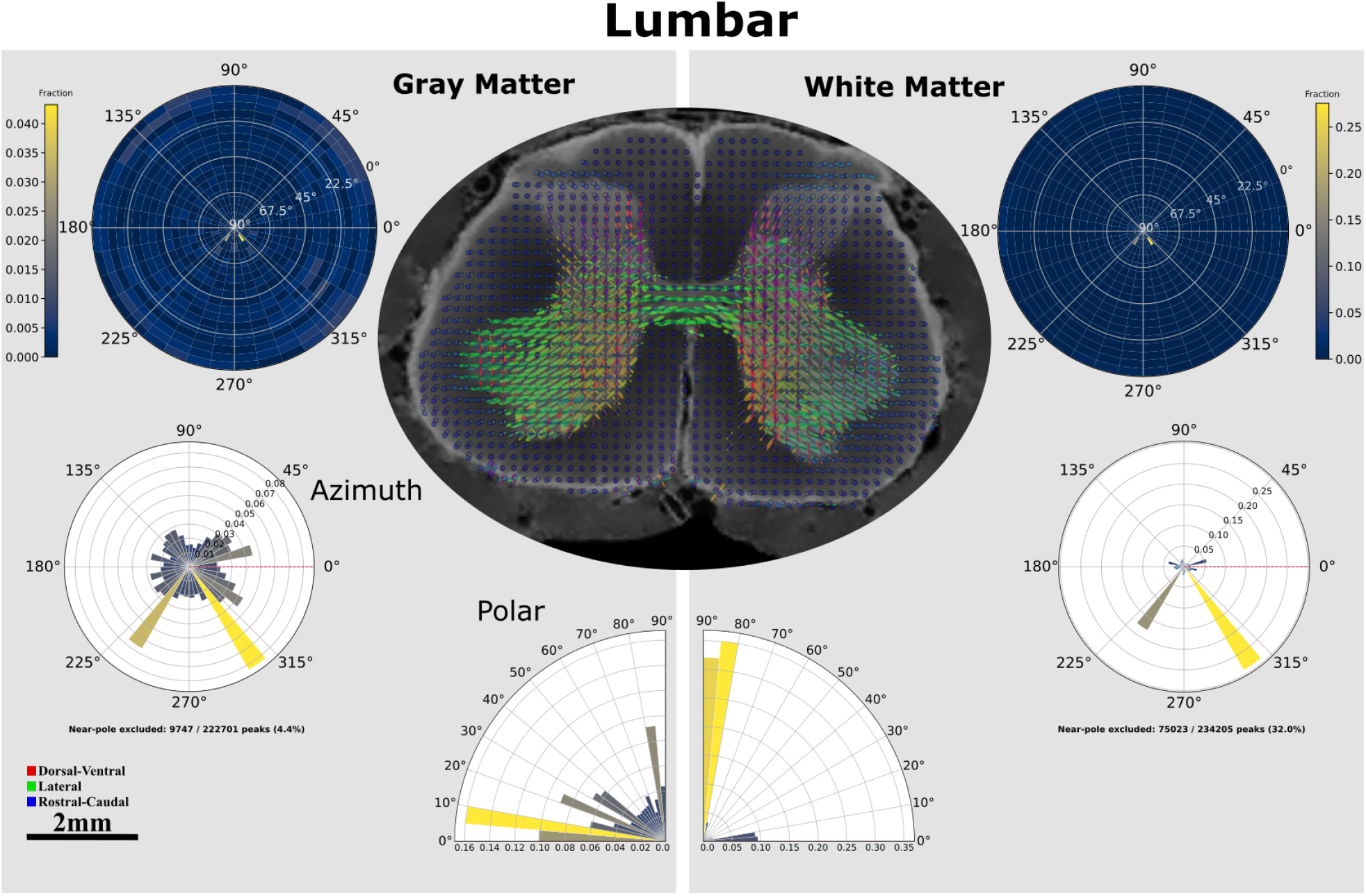
Quantifiation of lumbar fODF profiles on a hemisphere stratified by tissue type. Upper left and right depicts a density plot of a flattend hemisphere. With the radius i.e, the angles depicted in white denoting polar angles, and azimuth angles by the circular angles. Forming a fully 3d representation of the fODF rather than the individual polar and azimuth distributions shown below.

**Figure 16:**
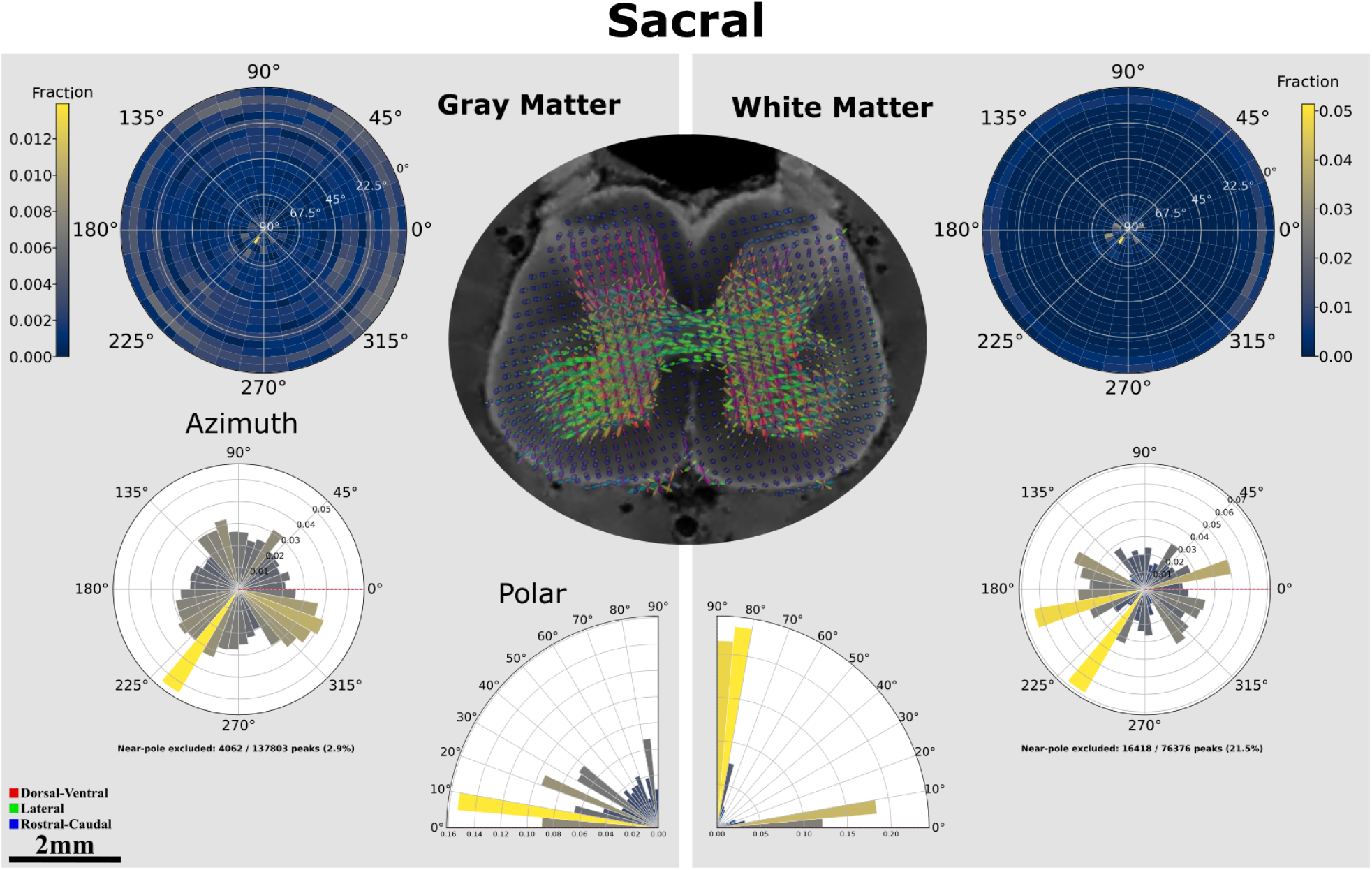
Quantifiation of sacral fODF profiles on a hemisphere stratified by tissue type. Upper left and right depicts a density plot of a flattend hemisphere. With the radius i.e, the angles depicted in white denoting polar angles, and azimuth angles by the circular angles. Forming a fully 3d representation of the fODF rather than the individual polar and azimuth distributions shown below.

**Figure 17:**
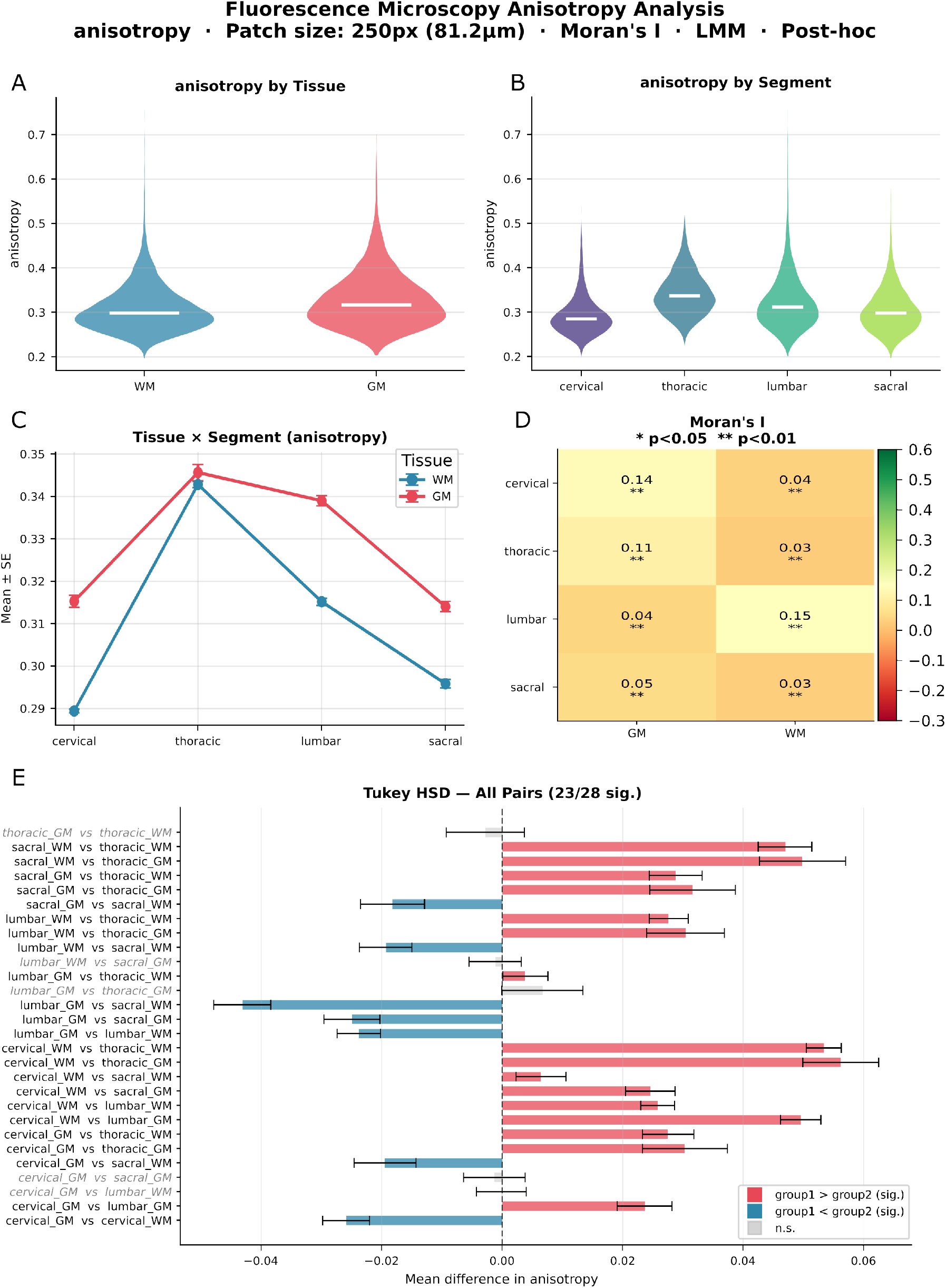
Overview of in depth statisical tests performed on structure tensor derived anisotropy from microscopy image. A & B) Shows the mean anisotropy across segment and tissue type. C) depicts ansisotroy stratified by tissue type, segment and their interaction. D) Moran’s I a measure of spatial autocorrelation. E) Depicts all post-hoc comparisons from the linear mixed effect model anisotropyT̃issue x Segment, with standard errors corrected for multiple comparison.

